# Representational similarity modulates neural and behavioral signatures of novelty

**DOI:** 10.1101/2024.05.01.592002

**Authors:** Sophia Becker, Alireza Modirshanechi, Wulfram Gerstner

**Affiliations:** School of Computer and Communication Sciences, EPFL; School of Life Sciences, EPFL; Helmholtz Munich; MPI for Biological Cybernetics

## Abstract

Novelty signals in the brain modulate learning and drive exploratory behaviors in humans and animals. While the perceived novelty of a stimulus is known to depend on previous experience, the effect of stimulus representations on novelty computation remains elusive. In particular, existing models of novelty computation fail to account for the effects of stimulus similarities that are abundant in naturalistic environments and tasks. Here, we present a unifying, biologically plausible model that captures how stimulus representations modulate novelty signals in the brain and influence novelty-driven learning and exploration. By applying our model to two publicly available data sets, we quantify and explain (i) how generalization across similar visual stimuli affects novelty responses in the mouse visual cortex, and (ii) how generalization across nearby locations impacts mouse exploration in an unfamiliar environment. Our model unifies and explains distinct neural and behavioral signatures of novelty, and enables theory-driven experiment design to investigate the neural mechanisms of novelty computation.

## Introduction

Novelty signaling in the brain is vital to facilitate learning [1–7], enhance sensory processing [8–12] and drive behavior [13–21]. Many experimental studies use computational models of novelty to explain and predict the effects of novelty signaling on neural activity and behavior [16–20, 22–24].

However, the concept of novelty and its quantification in these studies relies on the assumption that stimuli are either ‘identical’ or ‘different’. This approach ignores any further similarities between stimuli and causes substantial limitations in environments that are continuous or exhibit stimulus similarities. For example, consider two previously unobserved stimuli that are similar to each other but not identical, e.g. two paintings by the same artist that exhibit a similar painting technique and style. If the two stimuli are presented sequentially, then how ‘novel’ is the second painting after observing the first one? Classic novelty models force us to either (i) consider the two stimuli as separate entities, implying that the second painting is completely novel despite its similarity to the first painting that was just observed; or (ii) consider them as the ‘same’ stimulus, predicting that no matter whether we continue observing the first or the second painting, we get equally familiar with both of them. Both options are at odds with what we would expect from human and animal perception (see e.g. [25] for a review). In the same way, classic novelty models fail to address novelty computation in continuous environments, e.g. during spatial exploration. Indeed, several experiments suggest that the brain’s novelty response to a given stimulus does not merely depend on the number of times the exact same stimulus has been observed, but is also modulated by exposure to specific features of parts of the stimulus [13, 26–28] (see Discussion). This underlines the conceptual limitations of classic novelty models and raises the question how computational models of novelty can account for stimulus similarities.

In the field of machine learning, novelty-like signals are used to guide artificial agents during exploration of unfamiliar environments with sparse or no rewards [29–35]. These novelty-like signals are often computed as decreasing functions of the observed frequency of a given input (‘state’) to the artificial agent. The observed frequency is computed by neural networks that, through training, generalize across similar states [29–35]. The success of such novelty-guided artificial agents in exploring complex environments highlights the functional relevance of generalization across similar stimuli or similar locations in the environment. However, while machine-learning models of novelty computation deal well with continuous and complex environments, they rely on extensive training of separate networks to estimate novelty, as well as architectures and learning rules with limited biological plausibility and interpretability.

Here, we propose a unified modeling framework for novelty computation (‘similarity-based novelty’) that leverages probabilistic mixture models to combine the strengths of classic and ML-inspired novelty models. Our model is consistent with classic novelty models [16–20,22–24] (see [36] for a review) in environments with discrete and distinct stimuli, but it also extends to continuous spaces and environments with similarity structure. Our model implements novelty updates as a biologically interpretable plasticity rule that is consistent with various circuit models of novelty processing (e.g., based on adaptation, Hebbian synaptic plasticity or inhibitory circuits [37–40]) and allows to test experimental hypotheses about how novelty-computing circuits represent stimuli and stimulus similarities. We test our similarity-based novelty model in two data sets and derive new experimental predictions.

## Results

### Count-based novelty models fail to account for similarity modulation of novelty responses

Most algorithmic models of novelty computation in the brain [16–20, 22–24, 41, 42] are based on estimating the count-based frequency at which stimuli (e.g., sensory stimuli, spatial locations etc.) have been observed: The more often a stimulus is observed, the less novel it is. These ‘count-based’ novelty models usually consist of four building blocks: (i) a stimulus representation, (ii) a stimulus discretization, (iii) an empirical frequency function that measures the familiarity of stimuli, and (iv) a novelty function that computes the novelty of stimuli as a decreasing function of their familiarity [16–20, 22–24].

The stimulus representation specifies how stimuli from the environment are represented. For example, Gabor-like visual stimuli could be characterized by their angular orientation (Fig. 1 A, left). The stimulus discretization specifies how stimuli are counted. Count-based novelty models use separate counters for different stimuli. Every time a stimulus *s* is observed, its counter *C*(*s*) is increased by one unit (Fig. 1 A, middle). Examples where stimuli can be clearly identified as ‘distinct’ and ‘identical’ in this sense are human studies in abstract sequential decision-making tasks [17,43]. In continuous environments and for stimuli with continuous features like the orientation of Gabor filters, however, two stimuli are neither fully ‘distinct’ nor ‘identical’ – instead, they show varying degrees of similarity. In these settings, counting is not well-defined. Even though we can manually construct a discretization by binning the stimulus space, e.g. into equidistant bins based on psychometric discrimination thresholds, binning still fundamentally misses the continuous nature of stimulus similarities, and fails to capture their impact on novelty computation (see toy example below).

**Figure 1:**
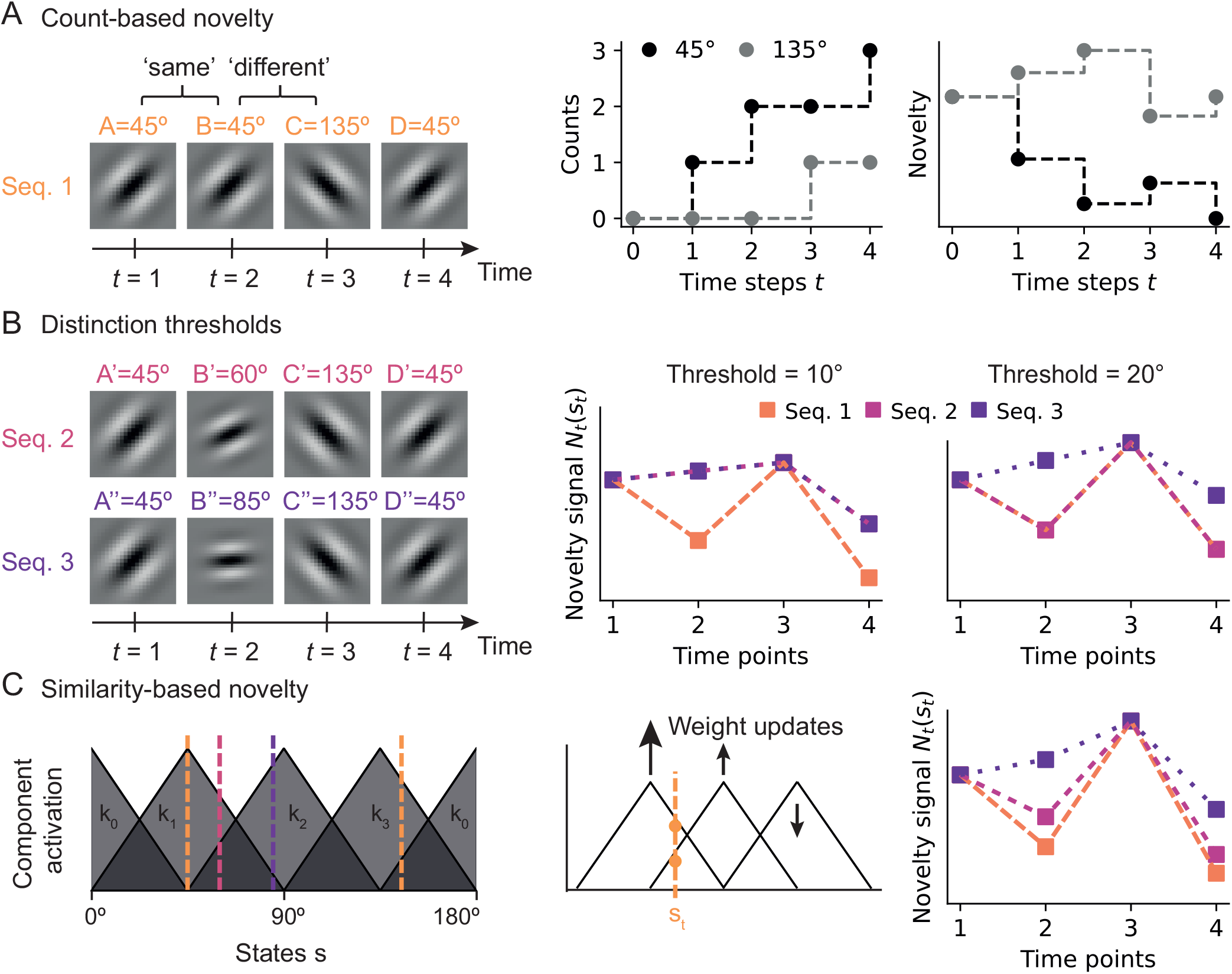
From count-based to similarity-based novelty. **(A)** Stimulus counts (middle) and count-based novelty responses (right) for an example sequence of four Gabors, characterized by their angular orientations between 0^°^ and 180^°^ (left). Each time we observe a stimulus, its count increases by one unit. Different stimuli (e.g. here: 45^°^ vs. 135^°^) are counted separately. Stimulus novelty is a decreasing function of the stimulus count. **(B)** Novelty responses for two count-based novelty models with different stimulus discretization. (left) Two additional example sequences of four Gabors each. The three sequences in (A) and (B) are identical except for the second stimulus, which shows varying level of similarity to the first stimulus in each sequence. (middle, right) Novelty predictions for the three Gabor sequences using two count-based novelty models. The models count stimuli as ‘different’ if they differ by more than 10^°^ (middle) or 20^°^ (right), respectively. The two count-based models make different predictions. **(C)** Similiary-based novelty using triangular components centered at reference orientations 0^°^, 45^°^, 90^°^ and 135^°^ (left). Component weights encode the empirical frequency of each component; they increase relative to the activation of their associated component (middle). Similarity-based novelty predictions (right) reflect the different levels of similarity between the first two stimuli across the three sequences: The novelty predicted for the second stimulus increases from seq. 1 to seq. 3, as the similarity between the first two stimuli decreases.

The third element of count-based novelty models is the empirical frequency function *p* that measures the familiarity of a given state *s* at time *t*:

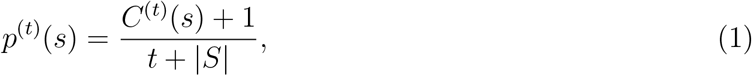

where *C*^(*t*)^(*s*) is the count of how often stimulus *s* has been observed up to the current time step *t*, and |*S*| is the number of different stimuli. At time *t* = 0, all stimuli start with the same prior frequency *p*^(0)^(*s*) = 1*/* |*S* |. The frequency increases if a stimulus *s* is observed (by increasing the count *C* and time *t*), and decreases if it is not observed (by increasing only the time *t*). In its most general form, count-based novelty allows the stimulus counts *C*^(*t*)^(*s*) to be ‘leaky’, i.e., decaying with a factor (1 − *α*) in each time step; *α* = 0 corresponds to the non-leaky count-based novelty model. Some count-based models compute the state familiarity using simpler functions than the stimulus frequency, e.g., absolute counts [16,22,41,44] or binary counts [18,23]. Since these models can all be reformulated in terms of the empirical frequency *p* [29, 36], we focus on frequency-based novelty models as in Eq. 1 for the rest of the manuscript.

The third element of count-based novelty models is the novelty function *N*. The novelty of a stimulus *s* decreases nonlinearly with its familiarity:

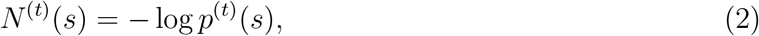

where the negative logarithm is a standard choice of nonlinearity in the novelty literature [36, 42].

The combination of empirical count-based frequency (Eq. 1) and logarithm (Eq. 2) makes the novelty decrease rapidly during the first few observations of a stimulus *s* (Fig. 1 A, right), in line with experimental observations [39].

To illustrate the behavior of count-based novelty models, we consider a toy example: We present three stimulus sequences with varying levels of stimulus similarities to two count-based novelty models with different state discretizations (Fig. 1 A-B, left). Each sequence consists of four Gabor stimuli that are each characterized by their angular orientation. The three sequences are identical except for the second stimulus, which has varying levels of similarity to the first (seq. 1: identical, seq. 2: 15° difference, seq. 3: 40° difference). We evaluate the novelty response to each stimulus as predicted by two count-based novelty models: The first model considers Gabors as distinct if they differ by more than 10° (Fig. 1 B, middle); the second one if they differ by more than 20° (Fig. 1 B, right). The two threshold values of 10° and 20° are consistent with psychometric thresholds for orientation differences measured in mice [45]. For the first sequence (yellow line in Fig. 1 B) and the third sequence (purple line in Fig. 1 B), both count-based novelty models predict qualitatively similar novelty signals: Every time we observe the first stimulus, its novelty decreases (seq. 1: at *t* = 2, *t* = 4, seq. 3: at *t* = 4), while previously unseen stimuli show high novelty responses (seq. 1: *t* = 1, *t* = 3, seq. 3: *t* = 1, *t* = 2, *t* = 3). Since we are measuring frequencies, and the initial counts are zero for all stimuli, the responses to completely novel stimuli increase during the observation of the sequence (with different slopes, depending on the discretization, i.e. the number of ‘bins’). Importantly, for the second sequence (pink line in Fig. 1 B), the two count-based models predict different novelty responses. The more fine-grained discretization (Fig. 1 B, middle) predicts the same novelty responses for seq. 2 and seq. 1 (‘identical’ first two stimuli), while the more coarse-grained discretization (Fig. 1 B, right) predicts the same responses to seq. 2 and seq. 3 (‘distinct’ first two stimuli). This exemplifies a fundamental limitation of count-based novelty models: They do not account for stimulus similarities beyond a simple distinction between ‘same’ and ‘different’. In naturalistic stimulus spaces, where stimuli vary continuously and stimulus similarities are abundant, these model predictions contradict both common-sense intuition and experimental evidence [13, 26–28] (see Discussion).

### Similarity-based novelty

To address the limitations of count-based novelty models, we propose a generalized novelty model (‘similarity-based novelty’) that accounts for the effect of stimulus similarities and allows to model novelty in continuous stimulus spaces. Its core difference to count-based novelty models is that, instead of computing the familiarity of stimuli as a function of their *discrete* empirical frequency (Eq. 1), our model uses probabilistic mixture models to compute a *continuous* empirical frequency density across the stimulus space *S*.

The idea of mixture models is to represent a complex probability density as the weighted sum of simpler probability densities (mixture ‘components’) over the stimulus space (see e.g. [46]). The mixture components can have arbitrary shapes, as long as they are non-negative and their integral over the stimulus space is one. In general, each component can overlap with an arbitrary number of other components. In the context of our problem, mixture components can serve as a flexible way to express stimulus similarities in both discrete and continuous spaces. For example, in our toy example, we can choose overlapping triangular components to account for orientation similarities (Fig. 1 C). Each component encodes the similarity of any given stimulus to a fixed reference stimulus, i.e., the orientation at the center of the triangular component. The pairwise similarity between two arbitrary stimuli can thus be described by the overlap in components they activate.

Based on the components, we define a mixture model *p* that generalizes the empirical frequency in Eq. 1. It quantifies the relative familiarity of a stimulus *s* at time *t* as

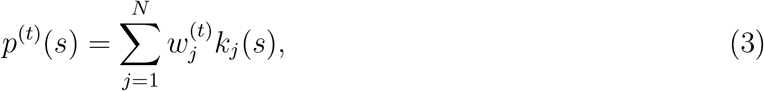

where *k*_1_, …, *k*_*N*_ are the components that account for the similarity structure of the stimulus space, and 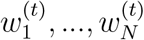, with 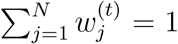, are the weights with which each component contributes to the empirical frequency density *p*^(*t*)^ at a given time point *t*. In order for the empirical frequency density *p* to represent a meaningful and consistent notion of familiarity across space and time, we must choose the weights 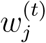 appropriately. We can do so using statistical inference: After a sequence of observations *s*_1_, …, *s*_*t*_, the ideal weights should be chosen so as to maximize the likelihood of the observed sequence. This normative approach leads to an iterative update rule:

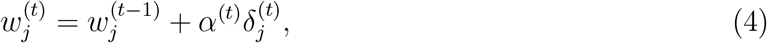

where 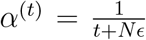 (see Methods) is the time-dependent learning rate, and the error 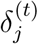 for the weight of component *k*_*j*_ at time *t* is

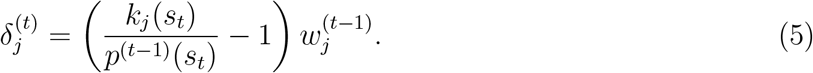

Crucially, the error in Eq. 5 depends on the previous weights through a multiplication factor, i.e., the familiarity weights are updated with a *multiplicative* learning rule, similar to what has been suggested in a range of mechanistic models of novelty detection [26, 37, 38, 47–49] (see Discussion). Since the error in Eq. 5 only relies on information that is locally available in time, i.e. the current component activation *k*_*j*_(*s*_*t*_) and the current familiarity *p*^(*t*−1)^(*s*_*t*_), it can be implemented in a biologically plausible neural circuit model (see Methods and SI). Analogously to the ‘leaky’ count-based novelty, we can define ‘leaky’ weight updates by choosing a constant weight update rate *α* in Eq. 4, resulting in an approximate maximum likelihood estimate of the component weights.

Using the learning rule in Eq. 4 along with suitable stimulus and component representations, we estimate the familiarity of a stimulus *s* at time *t* by its empirical frequency *p*^(*t*)^(*s*). Note that the definition of familiarity in Eq. 3-4 implies that similar stimuli influence each other’s familiarity via the components they share. For example, if a stimulus is covered by two (overlapping) components, its familiarity will be influenced by other stimuli that share any of the two mixture components. Analogously to count-based novelty (Eq. 2), we then apply the negative logarithm to *p*^(*t*)^(*s*) to compute the (continuous) novelty *N* ^(*t*)^(*s*). The resulting generalized novelty model (‘similarity-based novelty’) is (i) well-defined in discrete and continuous stimulus spaces, (ii) can account for similarities between stimuli via its component representation, and (iii) is equivalent to the established count-based novelty in discrete stimulus spaces without similarity structure (see SI).

To illustrate the differences between count-based and similarity-based novelty, we compare their novelty predictions in our toy example (Fig. 1 A-B, left). Analogous to the count-based model (Fig. 1 B, middle and right), we predict the novelty responses to each stimulus in each sequence using similarity-based novelty (Fig. 1 C). As components, we choose equidistantly placed, triangular functions that encode orientation similarities as linearly decreasing function of their angular difference (Fig. 1 C, left and middle). This resembles a linear approximation to the tuning curves of orientation-selective cells in V1 [50] (see [51] for a review). The predictions of similarity-based novelty are qualitatively different from the predictions of count-based novelty. In contrast to count-based novelty, similarity-based novelty predictions for the second stimulus increase with the angular difference to the first stimulus (Fig. 1 C, right), reflecting the varying degree of similarity between the first two stimuli across the three sequences. An analogous similarity-driven decrease of novelty responses is predicted for the stimulus at time *t* = 4. This shows that, in contrast to count-based novelty, similarity-based novelty generalizes familiarity across stimuli proportionally to their similarity, as defined by the underlying component representation.

### Similarity-based novelty explains novelty responses in mouse V1

In the previous section, we showed how we can model the impact of stimulus similarities on novelty computation using our similarity-based novelty model. In this section, we ask whether similarity-based novelty can capture key features of novelty responses in the brain, and provide further insights into how stimulus similarities modulate novelty computation in sensory areas. To this end, we compare the novelty signals predicted by count-based and similarity-based novelty with neural novelty responses in the mouse primary visual cortex (V1) during passive viewing of a sequence of ‘familiar’ and ‘novel’ images, as reported by Homann et al. [38].

The visual stimuli in the experiment of Homann et al. [38] are images that each consist of multiple randomly placed and randomly oriented Gabor filters (Fig. 2 A). Homann et al. study V1 novelty responses in mice during three variations of their passive viewing experiment (Fig. 2 C1-C2), each illustrating a specific qualitative feature of novelty responses in V1. In the first version of the experiment, a sequence of three images (*M* = 3) is presented for *L* repetitions (*L* = 1, 3, 8, 18, 38) before a ‘novel’ image (*X*) is presented in the subsequent repetition (Fig. 2 C1). Here, a ‘novel’ image is defined as an image that is not in the set of ‘familiar’ images that are shown as part of the repeating image sequence. The neural population response to the novel image increases with *L* (Fig. 2 D1, black line). In the second variation of the experiment, the number *M* of familiar images in the sequence is varied (*M* = 3, 6, 9, 12; Fig. 2 C1), while the number of sequence repetitions *L* remains constant (*L* = 18). Increasing *M* decreases the neural responses to the novel image (Fig. 2 D2, black line) but does not significantly affect the steady-state population activity after habituation to the familiar sequence (Fig. 2 D2, black line in inset). The third variation investigates the recovery from familiarity: after neural responses to a familiar sequence (‘adaptor’, *M* = 3) have converged, a novel sequence of images (‘replacement’, *M* = 3) is presented for *L*′ repetitions, before the initial sequence is shown again and the neural population response Δ*N*_recov_ is measured (Fig. 2 C2). The recovered novelty response to the formerly familiar sequence increases with *L*′ (Fig. 2 D3, black line).

**Figure 2:**
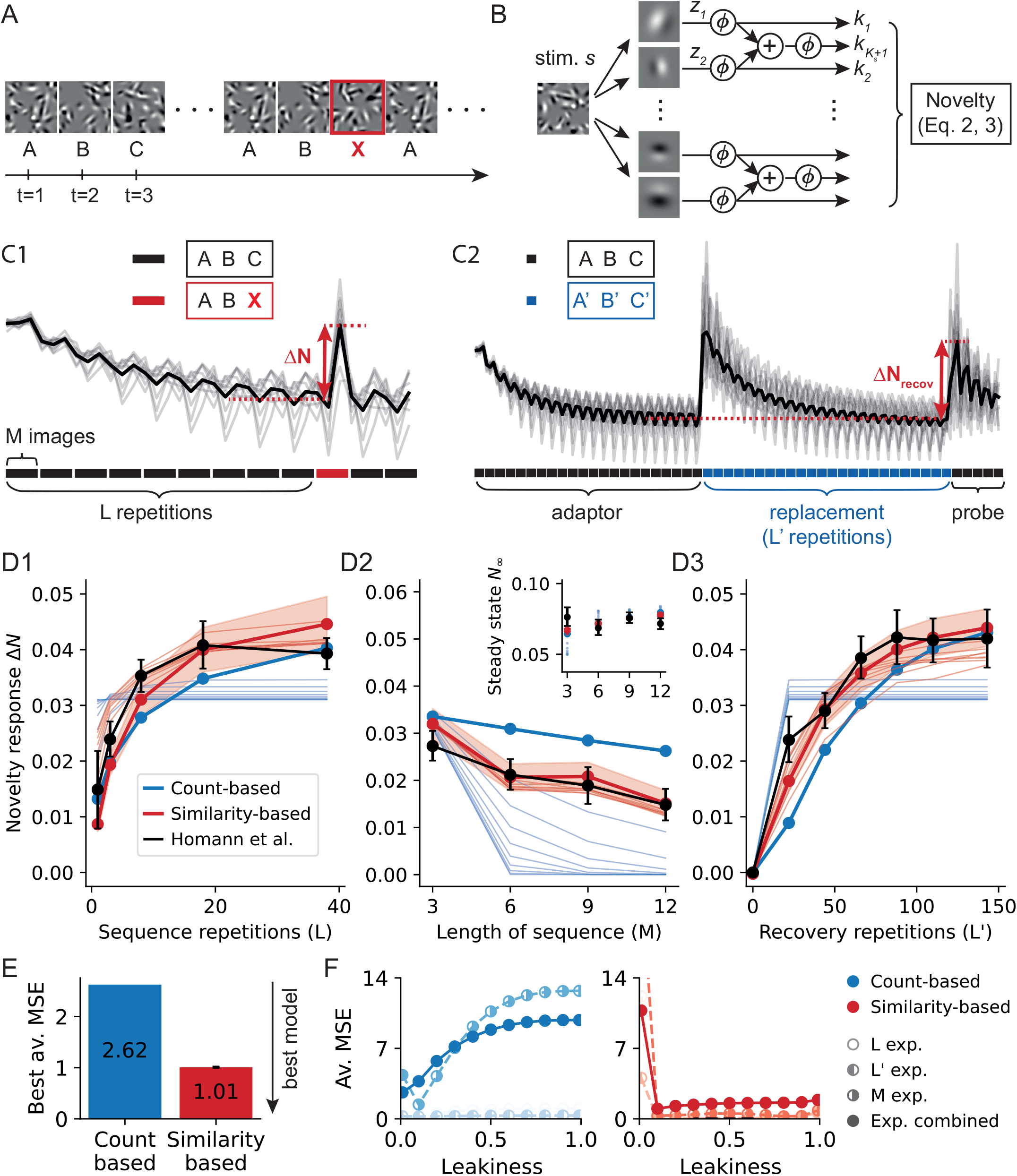
Similarity-based novelty robustly explains novelty responses in mouse V1. **(A)** Passive viewing task by Homann et al. [38]. A sequence of images is presented repeatedly, until the last image in the sequence is replaced by a novel, i.e. previously unseen, image (*X*). Familiar images in the sequence and novel images *X* are drawn from the same distribution and each consist of a linear superposition of 40 Gabor filters with random parameters. **(B)** The components of similarity-based novelty (*k*_*j*_ in Eq. 3) emulate V1 simple and complex cells [50, 51]. ‘Simple cell’-like components compute the cosine similarity between the stimulus image *s* and their reference Gabor (defined by its preferred orientation, phase, frequency, and location) and pass it through an arbitrary non-negative function *ϕ* (see Methods, here: ReLU, same for all components) to compute their component activation. The activation of a ‘complex cells’-like component is computed by applying the same function *ϕ* to the sum of 2-4 ‘simple cell’-like component activations with opposite phase and different frequencies. **(C)** Three variations of the passive viewing experiment in A. We consider responses to the ‘novel’ image for (i) different numbers *L* of sequence repetitions (*M* = 3 fixed; C1) and (ii) different numbers *M* of ‘familiar’ images in a sequence (*L* = 18 fixed; C1). We compare the novelty signals Δ*N* predicted by count-based and similarity-based novelty to the population responses of neurons in mouse V1 [38]. We further assess the recovery from familiarity in a third experiment (C2): after novelty responses to an initial sequence (‘adaptor’) have converged, a second, different sequence (‘replacement’) is presented for *L*′ repetitions; subsequently, the recovered novelty response Δ*N*_recov_ to the initial sequence (‘probe’) is measured (*M* = 3 fixed). **(D)** V1 novelty responses Δ*N* [38] (black line) and novelty predictions Δ*N* of fitted (best av. MSE) count-based (bold blue line) and similarity-based novelty models (bold red line) in all three experiments. For the *M* –experiment (D2), responses to the ‘novel’ image (Δ*N*) and steady state response to the ‘familiar’ sequence (*N*_∞_, inset) are shown. Thin blue and red lines depict novelty predictions of each novelty model for the remaining (non-optimal) values of the leakiness parameter *α* in F. Similarity-based novelty reproduces all qualitative features of neural novelty responses in V1, robustly for different values of *α*, whereas count-based novelty is highly susceptible to changes in *α*. **(E)** Average mean squared error (MSE) between neural novelty responses [38] and novelty responses predicted by count-based and similarity-based novelty, across the three experiments. **(F)** Average MSE between model predictions and neural responses across experiments as a function of the leakiness level *α* for count-based novelty (left, blue) and similarity-based novelty (right, red). Count-based novelty shows a bad fit (large av. MSE) everywhere except in a small region around leakiness *α* = 0.1 due to large errors in the M-experiment. Similarity-based novelty gives a robustly good fit (low av. MSE) across all leakiness levels (=novelty learning rates) except in the limit of very fast learning (*α* → 0). Average MSE is stable across other model parameters (see SI).

To test whether these qualitative features of novelty responses in V1 can be captured by count-based and similarity-based novelty, respectively, we fit each novelty model to the combined experimental data [38] from all three experimental variations (see Methods). The best-fitting parameters for each model are determined by minimizing the mean squared error (MSE) between the statistics of predicted and the experimentally measured neural novelty responses, averaged across the three experiments. For the count-based novelty model, all images are considered as distinct. For the similarity-based novelty model, we define components that emulate either V1 simple cells with specific orientation, phase, frequency and location preference, or V1 complex cells that combine the feature preferences of at least two V1 simple cells with opposite phase (Fig. 2 B, see Methods for details). The number of bins (for count-based novelty) and components (for similarity-based novelty) are free parameters that are fitted jointly with the remaining parameters of each model using grid search (see Methods).

In all three experiments, similarity-based novelty captures V1 novelty responses better than count-based novelty (Fig. 2 D1-D3 and E). Additionally, similarity-based novelty maintains its good fit across the majority of the parameter space, while the fit of count-based novelty is highly susceptible to parameter changes (Fig. 2 F, see SI for robustness results for remaining parameters). In particular, the fit of count-based novelty decreases rapidly with deviations from the fitted leakiness level *α* (Fig. 2 F, left panel). Similarity-based novelty captures the neural data well across the majority of leakiness levels *α* (Fig. 2 F, right panel). Only in the limit of very low leakiness levels *α*, the fit of similarity-based novelty decreases rapidly. This is to be expected since, for similarity-based novelty, the ‘leakiness’ level *α* is equivalent to the learning rate with which observations are integrated into the component weights; very small values of *α* thus prevent new observations from being incorporated into the familiarity estimate. The fit is stable with respect to the remaining model parameters for both novelty models (see SI). To further test the validity of our fit, we compare the statistics of novelty predicted by the fitted count-based and similarity-based model, respectively, with the statistics of neural novelty responses in V1 (Fig. 2 D1-D3). As expected, the novelty predicted by the similarity-based novelty model (red line in Fig. 2 D1-D3) captures the features of V1 novelty responses in all three experiments, robustly for fitted parameters (thick red line) and deviating leakiness levels (i.e., learning rates) *α* (thin red lines). In contrast, count-based novelty shows deviations from the V1 novelty responses in all three experiments (thick blue line) that rapidly increase as we deviate from the fitted leakiness level *α* (thin blue lines). Specifically, non-optimal count leakiness leads to instant saturation of the novelty responses predicted by count-based novelty in all three experiments (Fig. 2 D1-D3, thin blue lines).

Taken together, these findings suggest that count-based and similarity-based novelty reproduce V1 novelty responses using two qualitatively different mechanisms. Count-based trades off the timescales required for each experiment, resulting in high susceptibility to the leakiness parameter. For example, a higher than optimal count leakiness in the L-experiment causes the predicted novelty responses to reach their steady state in very few repetitions, preventing a gradual saturation of novelty responses as observed in the neural data. At the same time, a slower than optimal count-learning rate prevents the decline of novelty responses with increasing stimulus sequence length *M*. In contrast, similarity-based novelty reproduces V1 novelty responses using a largely generalization-based mechanism. For example, in the *M* –experiment, similarity-based generalization of familiarity leads to a natural decrease of novelty responses as the number *M* of familiar stimuli and, thus, the feature overlap with the novel stimulus increases. As a results, similarity-based novelty avoids a trade-off of learning timescales and ultimately achieves a better and more robust fit to neural responses across all three experiments.

### Similarity-based novelty explains mouse exploration in an unfamiliar maze

In the previous section, we showed that similarity-based novelty explains key features of novelty responses in mouse V1 during a passive viewing task [38]. In this section, we ask whether similarity-based novelty also helps explaining behavioral signatures of novelty processing during active exploration. To this end, we analyze behavioral data by Rosenberg et al. [52] who measure exploration of mice in an unfamiliar, freely accessible maze (Fig. 3 A). The maze has the structure of a binary tree (Fig. 3 C2): Mice enter the maze through a single corridor that then branches into two corridors, which each branch again into two corridors etc. After six branching points, each corridor ends in a wall (‘leaf node’), where mice can only turn around. Half of the mice do not receive any reward, the other half receive a drop of water upon licking a reward port in one of the maze’s leaf nodes (‘goal state’). To make sure that we only include novelty-driven behavior in our analysis, we consider mice behavior before the first encounter with the goal in both the unrewarded and the rewarded group. During this initial exploration phase, mice from the rewarded and the unrewarded group have access to identical information and do not show significant behavioral differences [52].

**Figure 3:**
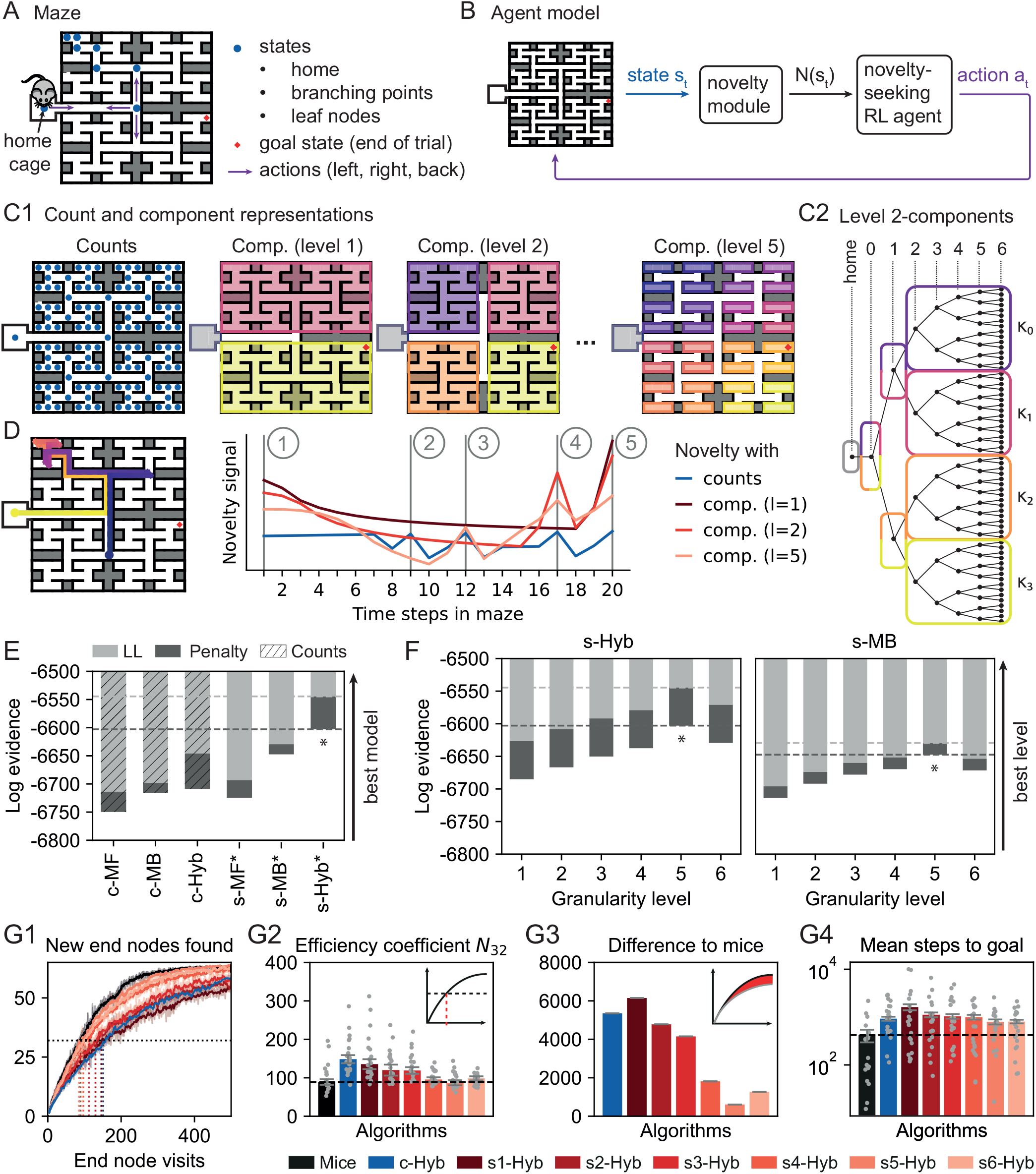
Similarity-based novelty explains mouse exploration in an unfamiliar maze. **(A)** The maze of Rosenberg et al. [52]. We define states (blue circles, example states on a short trajectory) with respective actions (purple arrows): (i) a single action in the home cage (go into the maze), (ii) three actions at ‘branching points’ (turn left, right or backwards), and (iii) one action in the ‘leaf nodes’ (turn back). To investigate intrinsically driven exploration, we consider mouse behavior before their first encounter with the rewarded ‘goal’ (red square). **(B)** In each time step *t*, a novelty-seeking RL agent takes its current position (state *s*_*t*_) as input and computes the state novelty using its internal novelty model. Then it updates its internal value function and policy based on a given RL algorithm (either model-free, model-based or hybrid), and chooses its next action *a*_*t*_ from its policy. **(C1-C2)** Counts and components used for the RL agents’ internal novelty models. (C1, first panel) State counts for count-based novelty. (C1, remaining panels) Components that tile the state space into ‘areas’ of the maze, e.g., into quadrants in the ‘level 2’ representation. Each component is active in a given area of the maze (marked in different colors) and in the states along the shortest path between the home cage and its encoding area. For example, the purple ‘level 2’ component *k*_0_ is active in all states within the purple quadrant and the two branching point states that connect it to the home cage (C2). **(D)** Novelty signals predicted by count-based and similarity-based novelty with different component granularity on an example path through the maze. Numbers indicate relevant behavioral events along the path: first entry into the maze (1), into novel level-6 component (2), level-5 component (3), level-2 component (4) and level-1 component (5). **(E)** Model fit of different novelty-seeking RL models to mouse exploration behavior. Log-evidence (loglikelihood augmented by penalty for number of parameters) of count-based (prefix: c-) and similarity-based (prefix: s-) novelty-seeking RL agents (model-free: MF, model-based: MB, hybrid: Hyb). For similarity-based novelty agents, the best model log-evidence across six granularity levels is reported (asterisk). Following the established interpretation scale for log-evidence [53, 54], a log-evidence difference of more than 3 is considered significant. **(F)** Comparison of similarity-based novelty with different granularity for the winning algorithm families (hybrid, model-based). Granularity 5 explains data best (asterisk). The best model overall is the hybrid RL agent with similarity-based novelty of granularity 5. **(G1-G4)** Posterior predictive checks of the model comparison in A-B. (G1) mean number of new leaf nodes found, as a function of total leaf node visits. Dashed vertical lines indicate the ‘search efficiency’ *N*_32_, i.e. the number of leaf node visits until 32 (half) of all 64 leaf nodes have been visited. (G2) Search efficiency *N*_32_); (G3) integrated difference between the end node discovery curves of agents and mice in G1; and (G4) mean number of steps until the first goal visit. Error bars indicate the standard error of the mean (sem).

We model mouse behavior using novelty-seeking reinforcement learning (RL) models that maximize intrinsically computed novelty instead of extrinsic rewards from the environment (Fig. 3 B, see Methods). In order to compare the exploration behavior of different ‘agents’, i.e. mice and novelty-seeking RL models, we define ‘states’ and ‘actions’ in the labyrinth. States include the home cage, the branching points and the leaf nodes (blue circles in Fig. 3 A). In each of the branching points, mice or RL agents can take one of three actions (purple arrows in Fig. 3 A): (i) going down the left corridor until the next branching point or leaf node, (ii) going down the right corridor until the next branching point or leaf node, or (iii) going back to the previous branching point. In the home cage and the leaf nodes, agents only have one available action, i.e. going back.

We compare mouse behavior to three families of novelty-seeking RL models, i.e., model-free (MF), model-based (MB) and hybrid (Hyb) RL algorithms (see Methods), and ask whether similarity-based novelty explains mouse behavior better than count-based novelty. For count-based novelty, we count each state in the maze separately. For similarity-based novelty, we define components that encode an abstract notion of spatial proximity in the binary tree by considering (i) how the mice got from the home cage to their current location (shortest path between a state *s* and the home state), and (ii) which ‘area’ of the maze the mice are in (e.g., a given side or a given quadrant of the labyrinth). The size of the ‘area’ encoded by a given component defines the spatial scale (‘granularity level’) across which it generalizes familiarity. For example, a ‘level 2’ component (Fig. 3 C1-C2) generalizes across all states in one quadrant and the shortest path leading to it from the home cage, while a ‘level 5’ component (Fig. 3 C1) generalizes across the two closest branching points of a given leaf node and the corresponding shortest path from the home cage. To test how this spatial scale of components affects novelty-driven exploration, we compare seven similarity-based novelty models with components of different granularity levels (levels 0-6, see Methods). Count-based and similarity-based novelty models with different granularity predict different novelty signals along an example path through the maze (Fig. 3 D).

To test whether exploration driven by similarity-based novelty helps explaining mouse behavior, we fit all of the model variants described above, i.e. all combinations of RL models (MF, MB, Hyb) and novelty models (count-based, similarity-based with different granularity) to mouse behavior, using maximum likelihood estimation of the parameters (see Methods). To determine which model family explains the data best, we then perform a Bayesian model comparison between the best-performing models of each family (see Methods). Across all types of RL models, exploration driven by similarity-based novelty gives a significantly better fit to mouse behavior than exploration driven by count-based novelty (Fig. 3 E). Model comparison between model-based and hybrid similarity-based agents shows that components of granularity 5, which generalize across two immediately neighboring leaf nodes and their shared branching point, best explain the exploration behavior of mice (Fig. 3 F).

To further validate our model comparison results, we perform posterior predictive checks [55] (see Methods). In particular, we simulate count-based and the different similarity-based novelty-driven RL models and compare them to mice, with respect to the efficiency of their exploration (Fig. 3 G). As expected, the winning model (similarity-based novelty-driven model with level-5 granularity) approximates mouse behavior best and shows the most efficient exploration efficiency among all tested models (Fig. 3 G). In contrast, both the absence of generalization (count-based novelty) and over-generalization (similarity-based novelty with broad components) show significantly lower exploration efficiency than mice and the winning model. Taken together, these findings suggest that spatial similarity structures significantly modulate exploration of mice in unfamiliar environments via their influence on novelty signals.

### Modeling novelty computation across naturalistic tasks

In this work, we propose a unified modeling framework for novelty (‘similarity-based novelty’) that generalizes the notion of novelty to continuous stimulus spaces and captures the effect of stimulus similarities on novelty (Fig. 4, left box). Its two main elements are (i) the set of non-negative component functions *k*_*j*_ that encode stimulus similarities in the representational space, and (ii) the history– and time-dependent component weights 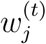 that encode the frequency of each component and are updated with a bio-plausible multiplicative learning rule (Eq. 4, 5). The familiarity of a given stimulus *s* is computed as the weighted sum of all component activations in response to *s* (Eq. 3), and the novelty of *s* is the negative log of its familiarity (Eq. 2). Since the weight update rule is directly derived from the normative approach and guarantees the frequency character of the familiarity density *p*, it remains the same across different choices of component sets. The component functions, on the other hand, can be flexibly adapted to model different data sets. While the similarity-based novelty framework allows arbitrary *non-negative* component functions, most component functions relevant for modeling consist of three elements: (i) a set of reference stimuli *r*_*j*_ that form ‘representative’ samples for similarity computation in the representational space of the input stimuli; (ii) a similarity measure *f* that is defined on the representational space; and (iii) a non-negative transform function *ϕ* that converts similarities into positive component activations.

**Figure 4:**
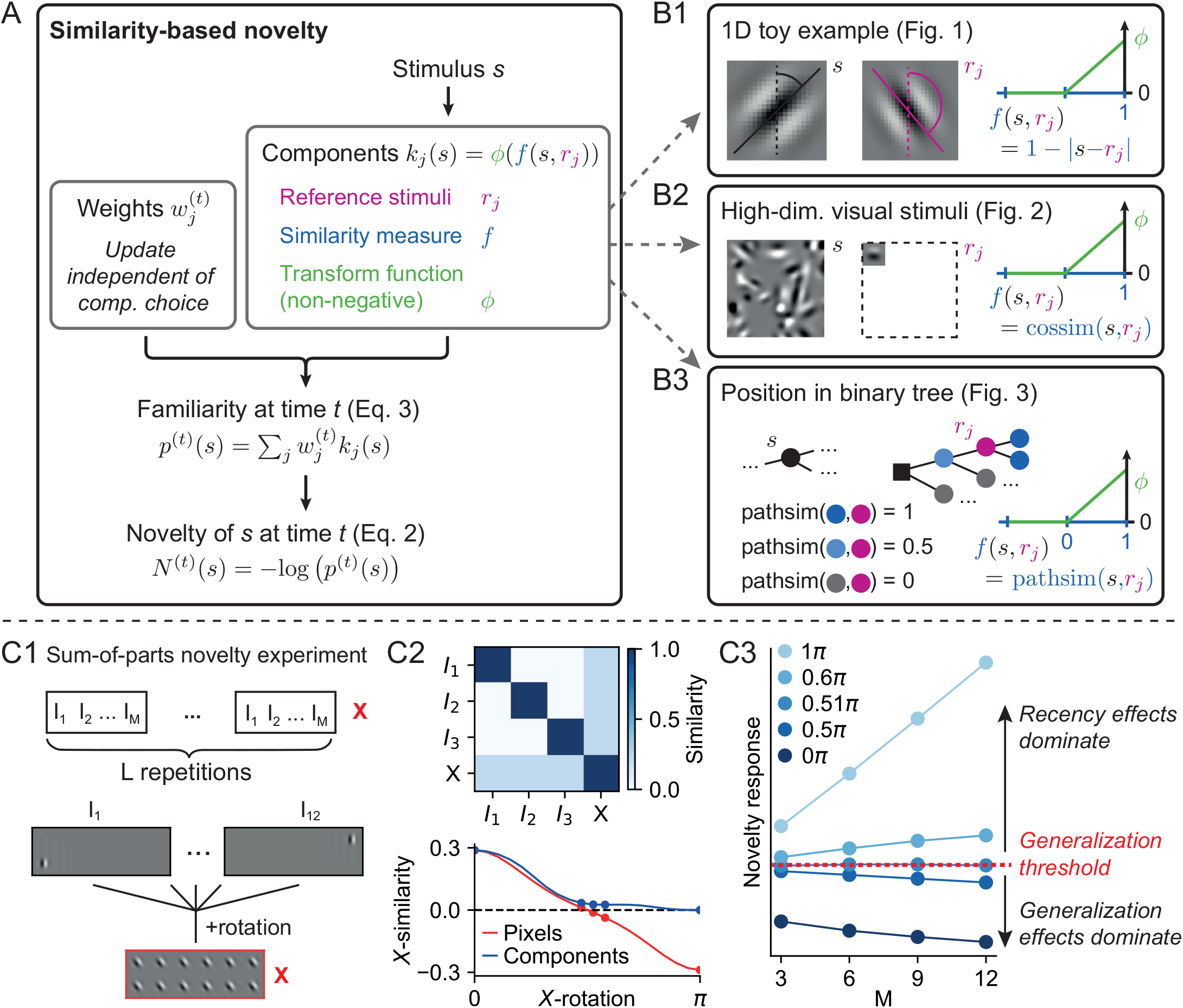
Modeling novelty computation across naturalistic tasks. **(A)** The similarity-based novelty framework represents the familiarity of stimuli as a probabilistic mixture model, using (i) non-negative component functions that capture similarities between stimuli, and (ii) component weights encoding the empirical frequency of each component in past observations. **(B1-B3)** The similarity-based novelty framework can be adapted to modeling neural data and behavior across different naturalistic tasks, enabling task– and stimulus-specific novelty predictions with a single probabilistic framework. **(C1-C3)** Schematic of a theory-inspired experimental protocol to further investigate the effects of stimulus similarities on novelty computation. (C1) A sequence of *M* familiar images (*M* = 3, 6, 9, 12) is presented for *L* repetitions, before a ‘novel’ image *X* is presented. Familiar stimuli are chosen such that they make up equal parts of the ‘novel’ image *X*. The degree of similarity between familiar sequence and the image *X* is varied by rotating the Gabors in *X* compared to the Gabors in the familiar sequence. (C2) Equal pairwise similarity between images in the familiar sequence (here: 0 similarity), and between each familiar image and *X* (upper panel). Higher rotation of *X* leads to lower pairwise similarity between any familiar image and *X*. Circles denote the rotation values for which novelty predictions were computed in C3. (C3) Novelty signals as predicted by similarity-based novelty for different values of *M* and different angles of rotation of *X* (i.e. different similarity between familiar and novel images). For high similarity (low rotation), novelty responses decrease with *M* (dark blue, ‘generalization’-dominant regime); for low similarity (high rotation), novelty responses increase with *M* (light blue, ‘recency’-dominant regime).

In our first toy example, we chose the reference stimuli *r*_*j*_ to be equidistantly spaced orientations in the 1-dimensional space of Gabor orientations, the similarity *f* to be measured as Euclidean distance of angles, and the transform function *ϕ* to be a triangular function that can be thought of as a simple approximation of Gaussian receptive fields (Fig. 4, right upper box). To model novelty for the more complex visual stimuli of Homann et al.’s experiment [38], we choose reference stimuli *r*_*j*_ that emulate the preferred features of simple and complex cells in V1, cosine similarity as a similarity measure *f* in the pixel space, and the same triangular transform function *ϕ* as in the toy example (Fig. 4, right middle box). Importantly, the reference stimuli in this application are defined in the same space as the input stimuli (pixel space) but – in contrast to the toy example – are not drawn from the same distribution as the input stimuli. To capture novelty in the binary tree maze of Rosenberg et al. [52], we defined branching points of the maze as reference stimuli *r*_*j*_, a shortest path-based spatial distance measure *f*, and the same triangular transform *ϕ* as in the previous two data sets (Fig. 4, right lower box). This exemplifies both the flexibility of the similarity measure and the possibility to combine our novelty framework with intrinsically motivated RL to model behavior. Thus, with a single probabilistic framework for novelty, we can model novelty computation, and its modulation by stimulus similarities, across passive and active tasks and behavioral and neural data sets.

Our findings suggest that stimulus similarities modulate novelty signals and novelty-driven behaviors in mice. However, while several studies [16, 21, 24, 38] test the impact of recency and repetition on the novelty of a stimulus, we still lack systematic studies about how stimulus similarities impact novelty computation. Using similarity-based novelty, we provide testable experimental predictions in a novel experimental protocol (‘sum-of-parts’ experiment, Fig. 4 C1-C3). Like in the *M* –experiment of Homann et al. [38], a sequence of familiar images consisting of *M* images (*M* = 3, 6, 9, 12) is presented for *L* repetitions, before a novel image *X* is presented. In contrast to the previous experiments, however, familiar stimuli are chosen such that they make up equal parts of the ‘novel’ image *X* (Fig. 4 C1). To vary the degree of similarity between familiar sequence and the image *X*, a controlled transform is applied to *X* (here: rotation of the Gabors in the image *X*). Importantly, the stimuli in the familiar sequence are fully distinct (zero pairwise similarity) but each of them has equal similarity to the novel image *X* (Fig. 4 C2, upper panel). The higher the rotation of the novel image *X*, the lower the similarity between familiar images in the sequence and *X* (Fig. 4 C2, lower panel). The same experiment could also be performed, e.g., with more naturalistic images as long as they preserve the similarity relationships between familiar and novel images. Using similarity-based novelty, we predict the novelty responses for different lengths *M* of the familiar sequence (*M* = 3, 6, 9, 12) and for different levels of similarity between the familiar and novel images (Fig. 4 C3). Similarity-based novelty predicts that, for high similarity between familiar and novel images, novelty responses decrease with *M* due to similarity-based generalization of familiarity. For low similarity between familiar and novel images, we predict the *opposite* trend, i.e. that novelty responses increase with *M* due to the increased number of total stimulus presentations in longer sequences. Our model simulations suggest that, as the similarity between familiar and novel image increases, the novelty responses gradually shift from the ‘generalization’-dominant regime (high similarity, dark blue in Fig. 4 C3) to the ‘recency’-dominant regime (low similarity, light blue in Fig. 4 C3). This example protocol illustrates how similarity-based novelty enables theory-driven experiment design.

## Discussion

We developed a generalized model of novelty computation that leverages mixture models [46] to capture the effect of stimulus similarities on novelty computation, and extends current computational novelty models to continuous state spaces. In tasks where all stimuli are discrete and distinct, our similarity-based novelty model is equivalent to count-based novelty [16–19,22–24,41,44]. We showed that similarity-based novelty captures novelty responses in mouse V1 [38] better and more robustly than count-based models. By combining similarity-based novelty with intrinsically motivated RL models [17, 22, 25, 29, 56], we further showed that seeking similarity-based novelty explains the exploration behavior of mice [52] better than seeking count-based novelty. Lastly, we use our similarity-based novelty model to make testable predictions in a novel experimental protocol.

Our results suggest that representational similarities between unfamiliar and familiar stimuli significantly influence both neural and behavioral signatures of novelty-related processing in the brain. Specifically, (i) similarity to familiar stimuli attenuates the novelty response to a previously unobserved stimulus, and (ii) generalization of familiarity across nearby spatial locations makes exploration more efficient by reducing the time spent exploring ‘similar’ (=close-by) regions in the maze. Our model unifies these findings by showing that they can both arise due to the modulation of novelty signals by stimulus similarities. Building on this, we predict that also novelty responses in other sensory modalities [4, 10, 27, 57] and in the hippocampus [6, 9, 58] as well as downstream novelty signaling in salience-related regions [5, 20, 21, 28, 59] could be subject to similarity modulation. If this prediction is true, it would have important implications for the experimental study of novelty: ‘novel’ input stimuli should either be chosen as almost perfectly distinct (e.g. [7, 17, 28]) to eliminate the impact of stimulus similarities on novelty signals, or their similarity should be controlled for as an experimental variable. Apart from enabling theory-driven experimental hypotheses and experiment design, our similarity-based novelty model also allows us to reinterpret existing experimental findings in the context of similarity-driven generalization of familiarity. For example, a study by Montgomery [13] investigating the impact of luminance on mice exploration in otherwise identical mazes finds that a larger luminosity difference (i.e. less similarity) between two successively explored mazes leads to more exploration in the second maze than a small or no luminosity difference (i.e. high similarity). This relationship is a signature of exploration driven by similarity-modulated novelty. Along the same lines, DeBaene and Vogels [26] report that neuronal adaptation in the monkey IT cortex significantly depends on the difference between adaptor and test stimulus, with respect to features such as object shape and location. This links directly to possible circuit implementations through which stimulus similarities could modulate novelty signals.

Moreover, insights from machine learning support the functional need to generalize novelty information across similar stimuli [25]. In particular, Jaegle et al. [25] argue that any system that computes novelty has to solve the problems of (i) mapping a diversity of high-dimensional inputs to reasonable states that abstract away e.g. different views on the same object (‘view invariance’), and (ii) grouping states together based on their shared characteristics (‘state invariance’). Similarity-based novelty solves the second problem by expressing similarities in a given representation space by (potentially overlapping) component functions. In our work, we addressed the first challenge, i.e., the choice of an appropriate state representation, by constructing stimulus representations based on experimental knowledge and hypotheses. In general, however, similarity-based novelty is not limited to such ‘experimentally justified’ representations and can also be used, e.g., in the latent space of trained deep networks with ‘brain-like’ representations [60,61]. As such, similarity-based novelty extends computational models of novelty to naturalistic tasks, allowing them to be modeled in a unified normative framework. Some machine learning algorithms solve both view and state invariance problem simultaneously by training a deep network to estimate a familiarity density from which novelty can be computed via pseudo-counts [29, 30]. Others use hashing [31] to map similar states to discrete bins that can then be used with count-based novelty (see [62] for an interesting neural implementation), or construct meaningful latent spaces where the novelty of a state is defined as a function of how many other states are within a given Euclidean distance in the latent space [34]. The fact that most of these methods are trained ‘end-to-end’ is both an advantage, since they learn an appropriate state representation simultaneously with the familiarity density, and a disadvantage, since they require learning of a separate density estimation network using backpropagation instead of computing novelty from existing representations in a modular fashion. In particular, they are not compatible with hand-designed representations based on ‘experimental knowledge’, making it harder to investigate, e.g., how a certain distribution of place fields influence novelty-driven exploration, or how novelty signals in V1 depend on the specific tuning of their cells.

One of the central advantages of similarity-based novelty is its biologically interpretable (local) learning rule. Specifically, the weights that capture the empirical frequency of each component are updated with a multiplicative learning rule. While similarity-based novelty does not make explicit predictions about which circuit structure has to underlie novelty computation, its update rule for the novelty weights is consistent with several types of circuit mechanisms proposed for novelty computation, including input adaptation and short-term synaptic depression [26, 37, 38, 47], inhibitory circuits [40], or Hebbian/anti-Hebbian multiplicative plasticity [48, 49]. As such, similarity-based novelty proposes a normative perspective on a wide range of mechanistic novelty models that provide area– and modality-specific mechanistic implementations of novelty computation.

While novelty has been shown to be a dominant drive of human exploration in unfamiliar environments [17, 19], a diversity of other intrinsic reward-like signals are involved in driving exploration [9, 13–15, 17–19, 21, 63, 64] (see [59, 65–68] for reviews), especially when extrinsic rewards such as food and money are absent. The relative importance of different motivational signals in explaining behavior depends on multiple factors, including the temporal stages of exploration (see e.g. [17]) as well as the exploration objectives and features of the task and environment (see e.g. [69]). Our findings for novelty suggest that stimulus similarities might be another aspect that substantially influences the computation of these intrinsic signals and their impact on behavior. In neuroscience, however, most of these motivational signals are investigated in a count-based setting. The question how stimulus similarities impact the computation of these diverse set of motivational signals, including surprise and information gain, thus remains an important direction for future research.

In conclusion, we propose a unifying computational framework to model novelty computation in the brain (‘similarity-based novelty’) that allows to assess the impact of similarity-driven generalization on neural novelty responses and novelty-driven behavior. Our approach opens new possibilities for future theory-driven experiment design.

## Methods

### Leaky updates for count-based novelty

In its most general form, count-based novelty (Eq. 1, 2) allows for ‘leaky’ updates of the stimulus counts *C*^(*t*)^(*s*):

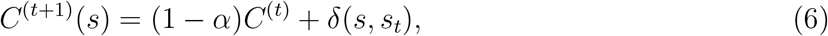

where *α* is the decay parameter, and *δ* is the Kronecker delta function (1 if *s* = *s*_*t*_, 0 elsewhere). The larger *α*, the faster the counts decay over time. For *α* = 0, we recover the ‘non-leaky’ classical count-based novelty model.

### Normative update rule for similarity-based novelty

The central element of similarity-based novelty is the mixture model that defines the familiarity density *p* (Eq. 3). Its weights at time point *t* are chosen as the maximum-a-posteriori (MAP) estimate of the sequence *s*_1:*t*_ of stimulus observations up to *t*:

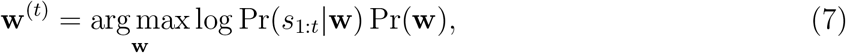

where 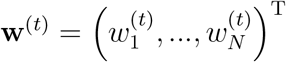 is the vector of mixture weights at time *t*, and Pr(**w**) is a Dirichlet prior over the weights (see SI). We solve Eq. 7 using the incremental expectation-maximization (EM) algorithm [46, 70, 71]. The resulting closed-form solution for the weights can then be reformulated as an online learning rule given in the main text (Eq. 4). The two weight update rules (batch and online) derived from the EM algorithms are given as follows.

i. **EM update (batch)**. For each time point *t* at which we observe a stimulus *s*_*t*_, we compute the E-step (estimation step) of the incremental EM algorithm, i.e. we compute ‘responsibilities’ *γ*_*j*,*t*_ that estimate how well a given component *k*_*j*_ of the mixture distribution at time *t*− 1 captures the observation *s*_*t*_:

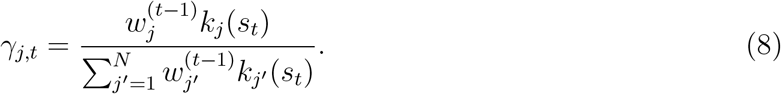 The initial weights are chosen uniformly as 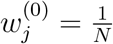 for all *j* = 1, …, *N*. In the subsequent M-step (maximization step) for time point *t*, we compute the new weights 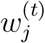 based on the responsibilities 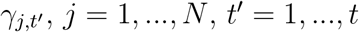 that were estimated *in all previous* E-steps (Eq. 8):

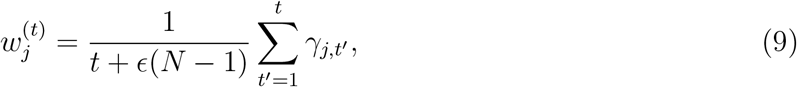 where *ϵ* is the uniform concentration parameter of the Dirichlet prior over the weights. The main difference to textbook incremental EM algorithms [46] is that at each iteration, the amount of data in the batch increases since an additional data point has been added to the sequence *s*_1_: *t* (see SI for details).
ii. **Online update**. The expression for the weights at time *t* (Eq. 9) still depend on the information from previous E-steps through the ‘responsibilities’ 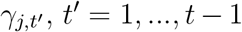 However, we can reformulate Eq. 9 to obtain an online weight update rule that computes **w**^(*t*)^ only based on the responsibilities from the current E-step at time *t* and the previous weights **w**^(*t*−1)^:

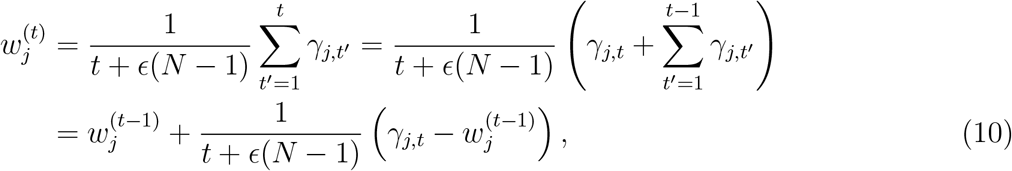 for all *j* = 1, …, *N*. The learning rule in Eq. 10 can be written as a delta learning rule as described in the main text (Eq. 4):

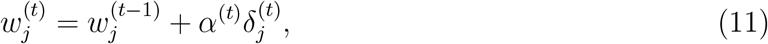 with error

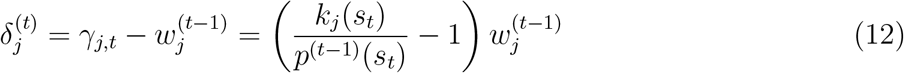 and either time-dependent learning rate (derived from incremental EM)

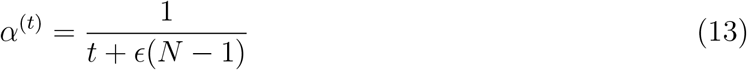 or constant learning rate (‘approximate’ incremental EM). A constant learning rate yields a ‘leaky’ weight update, analogously to the ‘leaky’ count-based novelty. Our learning rule in Eq. 11 is *multiplicative* since the error (Eq. 12) depends on the previous weights 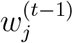 through a multiplication factor. The multiplication factor of the weights in Eq. 10 depends on the ratio of component activation and familiarity of the current observation *s*_*t*_, which are both available locally in time. For example, the component activation by the current observation *s*_*t*_ can be interpreted as the presynaptic activity of weight 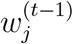, while the familiarity of *s*_*t*_ can be seen as the postsynaptic activity of the novelty-detecting circuit (see SI for a biologically plausible circuit implementation).

### Toy experiment: Gabors sequences of different similarity

Novelty predictions of count-based and one similarity-based novelty models were computed for three sequences of four Gabor stimuli each. Gabor stimuli are characterized by their angular orientation. Across all sequences *i* = 1, 2, 3, all except the second stimulus *s*_2_ are identical. The second stimulus in each sequence shows varying levels of similarity to the first stimulus: 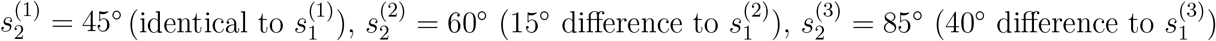.

Countable states for the count-based novelty models are obtained by binning the space of Gabor orientations [0, 180^°^] into 9 bins of width 20^°^ with centers 0^°^, 20^°^, 40^°^, etc. (model variant 1) or into four bins of 45^°^ with centers 0^°^, 45^°^, 90^°^ and 135^°^ (model variant 2). The size of the state space, |*S*|, is equal to the number of bins, respectively; counts at *t* = 0 were initialized at zero, and leakiness was set to *α* = 1 (non-leaky).

The component representation of similarity-based novelty is defined on the 1D torus of Gabor orientations [0, 180^°^]. We use four equidistantly placed components *k*_*j*_, *j* = 1, …, 4, centered at *c*_1_ = 0^°^, *c*_2_ = 45^°^, *c*_3_ = 90^°^ and *c*_4_ = 135^°^. The components are defined by the following triangular function:

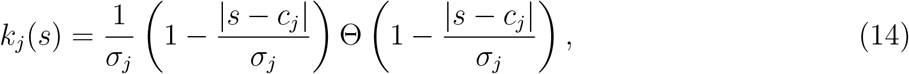

where *s* ∈ [0^°^, 180^°^] is the orientation of the Gabor stimulus, *c*_*j*_ as defined above are the centers of the components, and *σ*_*j*_ are the component widths. We choose *σ*_*j*_ = 90^°^ for all components, creating the overlapping components depicted in Fig. 1 C. The component weights at *t* = 0 were initialized uniformly to 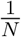, where *N* = 4 is the number of components. For the toy example simulations, we used non-leaky similarity-based novelty, with prior *ϵ* = 1.

### Passive viewing task: Experimental data by Homann et al

#### Setup and recordings

Homann et al. [38] used two-photon calcium imaging to record the neural activity of layer 2/3 neurons in the primary visual cortex (V1) of 5 GCaMP6f-expressing mice during passive viewing of ‘familiar’ and ‘novel’ visual stimuli. During the recording, mice were head-fixed but could run freely on an air suspended Styrofoam ball in front of a toroidal screen, onto which the visual stimuli were projected. Homann et al. extracted the peak neural responses (trial– and population-averaged, see SI for details) to the novel image (Δ*N*) and to the familiar sequence (steady-state activity *N*_∞_) during the ‘variable repetition experiment’ (*L*-experiment) and the ‘variable image number experiment’ (*M* –experiment), as well as the transient population response (Δ*N*_recov_) to the formerly familiar sequence after the recovery interval in the ‘repeated image set experiment’ (*L*′-experiment).

### Passive viewing task: Model simulations and fitting

#### Stimuli

All images for our simulation of the passive viewing experiment were sampled from a distribution, designed to match that of Homann et al. [38]. Each image is the linear superposition of 40 Gabor filters with randomly chosen orientation (uniformly sampled from [0, *π*]), phase (unif. sampled from [− *π, π*]), frequency (unif. sampled from [0.04, 0.08]), and location (*x* and *y* coordinate sampled uniformly from [− 130, 130] and [− 20, 90], in units of visual degrees) on a grey background. The width of each Gabor filter was set to 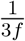, where *f* is the spatial frequency of the Gabor filter.

#### Simulation protocol

Model simulations followed the same protocol as the mouse experiment by Homann et al. Each simulation step corresponds to 300 ms in the experiment, i.e. in each simulation step, a single stimulus is presented.

#### Component definition

The input stimuli for similarity-based novelty are 100×200 pixel images as defined above. Each image is represented by a pixel vector *s* ∈ ℝ^20,000^ with entries specifying the intensity of each image pixel (grey-scale; the lower the pixel value, the darker the color; grey background has pixel value 0). We define two types of component functions that emulate the receptive fields of V1 simple and complex cells, respectively.

##### ‘Simple cell’-like components

Analogously to the component centers in the toy example, each component function *k*_*j*_ is defined relative to a ‘reference Gabor filter’ *r*_*j*_ ∈ ℝ^20,000^, i.e. the pixel vector of a single Gabor filter that is chosen equidistantly from the same parameter space (orientation, phase, frequency, *x*−*y*-location) as the Gabor filters that form the stimulus images. The value of component *k*_*j*_ with reference Gabor *r*_*j*_ in response to a stimulus image *s* is defined as

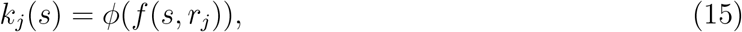

where *f* is the cosine distance between the pixel vectors *s* and *r*_*j*_:

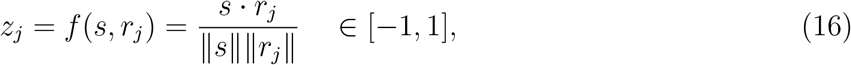

and *ϕ* is a non-negative transform function that maps the similarity value between the stimulus and the reference Gabor (in the interval [−1, 1]) to the component activation. We choose the function *ϕ* to be triangular with width parameter *σ*_*j*_ = 1 and centered at *z* = 1 (maximum similarity):

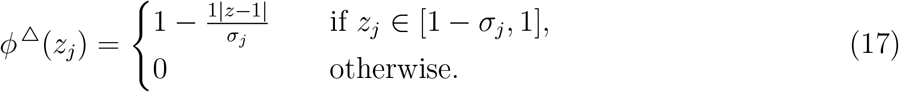

##### ‘Complex cell’-like components

‘Complex cell’-like components are defined relative to a reference set of two ‘simple cell’-like kernels of opposite phase. The activations from the set of reference components are added, and then passed through the same function *ϕ* as for the ‘simple cell’-like components. ‘Complex cell’-like components can be thought of as encoding an OR relation between the reference Gabors of the ‘simple cell’-like components of their reference set.

We define *K*_*s*_ ‘simple cell’-like components and *K*_*c*_ ‘complex cell’-like components. We keep the ratio of 3:1 of simple cell-like to complex cell-like components, in alignment with experimental results for layer 2/3 of V1 in mice [72], where Homann et al. recorded. The total number of components *N* = *K*_*s*_ + *K*_*c*_, the width *σ*_*j*_ = *σ* (shared across all components) and learning rate *α* that determines the speed of similarity-based novelty updates are free parameters that are fitted to the neural data by Homann et al.

#### Comparison with count-based novelty

The count-based model receives the same inputs but does not use receptive fields. Instead, separate counts are used for different images. The parameter *α* is fit to data.

#### Fit to neural data

To fit each novelty model to the neural data, we use grid search in combination with linear regression. We simulate a given novelty model ℳ ∈{count-based novelty, similarity-based novelty} across the full range of model parameters (*θ*_count_ = (|*S*|, *α*), *θ*_similarity_ = (*N, σ, α*), see SI); we then fit the simulated novelty responses to the neural data using linear regression (see SI). The parameter set that minimizes the normalized mean-squared error (MSE), averaged across the three experiments, is chosen as the best-fit parameter set for model ℳ.

### Active exploration task

#### Task

Rosenberg et al. [52] track the behaviour of 20 mice (10 of them water-deprived) during free exploration of an unfamiliar maze (maze cleaned with ethanol between mice). Each mouse was video-recorded during its first 7 hours of access to the maze The maze was constructed as a 6-level binary tree, with branching points and end points of corridors as nodes. One single end node in the maze contained a water delivery port, which delivered a water reward to water-deprived mice (‘rewarded group’) but was inactive during the experiment with non-deprived mice (‘unrewarded group’). The *x*-*y*-coordinates of the animal’s nose in each video frame were used as the ‘mouse position’ for subsequent behavioral analysis (both by Rosenberg et al. and in our study, see SI for details).

#### Environment formalization

To enable direct comparison between the behavior of mice and reinforcement learning (RL) models, we formalize the labyrinth environment using ‘states’ and ‘actions’. ‘States’ *s* characterize the possible locations of agents (mice or RL models) in the labyrinth maze, and ‘actions’ *a* characterize the possible next direction the agent can take from a given state. Analogously to Rosenberg et al. [52], the home cage and all branching points and end points of the labyrinths corridors (i.e. all nodes in the binary tree) were defined as ‘states’. We define four different actions: (i) go forward into maze (available in the home cage); (ii) go back (available in all states except the home cage); (iii) turn and go into the left corridor (available in all branching points) and (iv) turn and go into the right corridor (available in all branching points). For simplicity of notation, our notion of left and right is based on the egocentric view of the agent during an outward path. Based on the definition of states in the maze, mouse behavior over time is discretized into ‘time points’, where each time point marks the transition to a new state in the maze.

#### Preprocessing

For each mouse in Rosenberg et al.’s data set (github: [73]), we extract the sequence of states and actions taken at each time point between their first entry into the maze and their first encounter with the goal state. To exclude any reward-related behaviors from the analysis, we disregard mouse behavior after the first encounter with the goal state, independently of whether the mouse received a reward in the goal state (rewarded group) or not (unrewarded group). Since there are no significant behavioral differences between rewarded and unrewarded mice during the time before the first reward encounter [52], we treat them as equivalent (see SI for details on outliers).

### Novelty-seeking reinforcement learning (N-RL) models

We adapt classical reinforcement learning (RL) algorithms to model novelty-seeking (‘novelty-seeking RL (N-RL) agents’) by replacing the extrinsic reward signal by an intrinsically computed novelty signal (see e.g. [17, 22, 25, 29, 56] for similar approaches).

Like classical RL agents, N-RL agents move in an environment which is defined by states *s*, and actions *a* that allow the agent to transition between states. In general, the environment can be probabilistic in the sense that a given action *a* in a given state *s* can lead the agent into different states *s*′. The probabilities for each such state transition (*s, a, s*′) are called transition probabilities and summarized in the transition matrix *p* ∈ ℝ^|*S*|×|*A*|×|*S*|^, where |*S*| is the size of the state space *S*, and *A* is the size of the action space |*A*|. In our model, the environment is deterministic, i.e., the correct transition matrix contains only binary entries.

The main difference between novelty-seeking and classical RL algorithms is that, instead of maximizing the discounted future rewards, N-RL algorithms maximize the discounted future novelty:

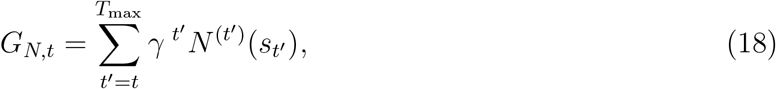

where *γ* is the discount factor, and the novelty 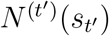 is computed using any suitable novelty model, e.g. count-based or similarity-based novelty. Analogous to reward-based V-values and Q-values, N-RL algorithms compute ‘novelty-based’ V-values and Q-values:

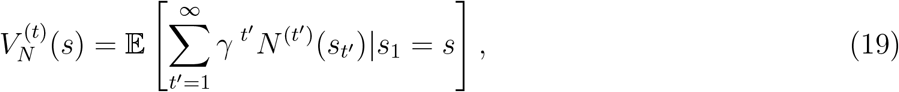

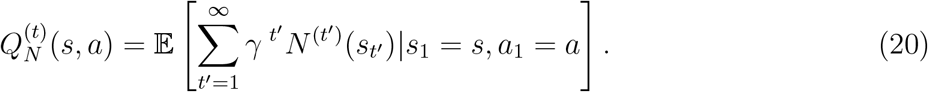

### Active exploration task: Model simulations and fitting

We simulate and fit a total of 24 novelty-seeking RL (N-RL) models to the exploration behavior of the mice in the Rosenberg maze, and compare the fitted models with respect to their log-evidence. We consider models from three different RL model families: (i) model-based N-RL models based on Prioritized Sweeping [74] (Alg. 1 in SI), (ii) model-free N-RL models with Actor-Critic architecture [74] (Alg. 3 in SI), and (iii) hybrid N-RL models that combines the model-free and model-based variants using a hybrid action policy (Alg. 4 in SI). For each N-RL model family, we consider one model with count-based novelty, and seven models with similarity-based novelty, one for each of the component representations (see below).

#### Count-based novelty

Count-based novelty in the Rosenberg maze uses a separate count for each state as defined above, i.e. each node in the binary tree representation of the maze. State counts are updated in a non-leaky fashion (*α* = 1). Using Eqs. 1 and 2, the novelty reward at each time is then computed as the novelty of the current state in the environment.

#### Similarity-based novelty

We define similarity-based novelty in the Rosenberg maze with respect to seven component representations that differ in the degree of granularity with which they encode the maze.

The component representations are based on the idea that each component should encode a different ‘area’ of the labyrinth environment, e.g. the left or right half of the maze (allocentric perspective), as well as a trace of how to reach this ‘area’ from the home cage. The size of the area encoded by the components determines the ‘granularity’ of the representation, i.e. the larger the area encoded by each component, the lower the granularity of the representation. To formally write the corresponding component functions, we consider the binary-tree structure of the labyrinth maze (Fig. 3C2) and define the ‘(sub-)tree’ Tree(*s*) for any state *s* in the labyrinth (*s* ∈ *S*) as

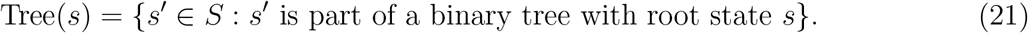

Hence, the entire labyrinth (excluding the home state) can be written as Tree(*s*_0_) where *s*_0_ is the first branching point of the labyrinth. We further define the sets *S*_𝓁_, 𝓁 = {0, 1, …, 6}, where each set *S*_𝓁_ contains all 2^𝓁^ states in a given level 𝓁 of the binary tree. For example, the set *S*_0_ contains only *s*_0_, the first branching point of the maze, while *S*_6_ contains all 64 leaf nodes.

We define the component representation of granularity level 𝓁, for 𝓁 = 1, …, 6 as follows. The component representation of a given granularity 𝓁 consists of 2^𝓁^ + 1 components: first, the home cage component, that is the same across all granularity levels and encodes the home cage separately:

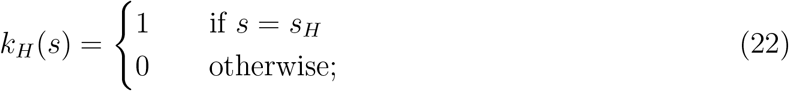

and second, the components 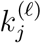 that each encode the ‘area’ around one of the states 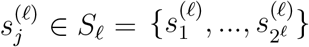

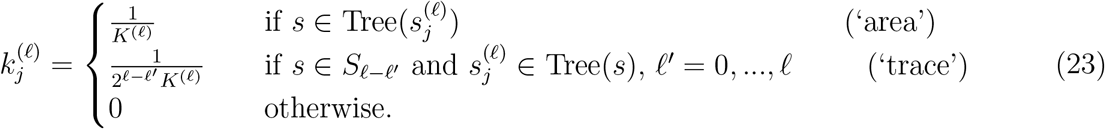

The factor 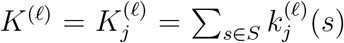 is added to normalize the components to 1 over the state space.

#### Model fitting

We fit the parameters of each N-RL model to mouse behavior using maximum-likelihood estimation (MLE):

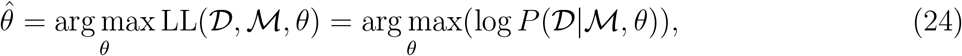

where ℳ is the N-RL model that is being fitted, *θ* are its model parameters, and 𝒟 is the mouse data. The mouse data 𝒟 consists of the appended state-action sequences of all mice during their exploration of the labyrinth:

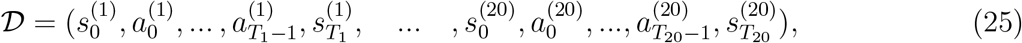

where 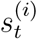 and 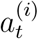 denote the state and action of mouse *i* at time point *t*, and *T*_*i*_ denotes the number of exploration steps of mouse *i* until its first encounter of the goal state.

We maximize the log-likelihood for a given model by minimizing its negative log-likelihood using the scipy.optimize implementation of the Nelder-Mead algorithm (best performing among relevant optimization algorithms, including L-BFGS-B, SLSQP). In each minimization step, the log-likelihood for the current parameter set *θ* is computed (see SI), and its negative is used as input to the minimizer of the scipy.optimize package to compute the next candidate parameter set. We run the optimization until convergence to obtain the fitted parameters for a given model.

#### Bayesian model comparison

We compare fitted models with respect to their their log-evidence (LE):

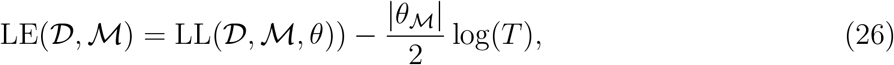

where the log-likelihood term accounts for the fit of the model, and the remaining term introduces a cost for model complexity, that is a function of the number of free parameters |*θ* _ℳ_| of the model, and the number of data points 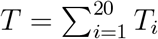 in the fitted data set 𝒟,

#### Posterior predictive checks

We perform posterior predictive checks to verify that the winning model from our model comparison matches the fitted data (i.e. mouse behavior) with respect to relevant statistics of their behavior. To this end, we simulated all fitted models (*n* = 500 instantiations for each model). Each model simulation starts with the model agent being initialized in the home state, and ends when the model agent first reaches the goal state.

We compare mice and simulated models with respect to (i) the average number of steps in the maze until reaching the goal state; (ii) the average number of novel end nodes visited relative to the total number of visits to a (familiar or novel) end node; (iii) the average search efficiency *N*_32_, i.e. the total number of end node visits until 32 (i.e. half) of the end nodes in the maze have been discovered; (iv) the integrated difference between the average end nodes discovery curve of mice and the average end node discovery curve of a given model. For the fourth statistic, the sem is computed using bootstrapping on the set of end node discovery curves of individual agents of a given type.

## Supplementary Information

### 1 Novelty models

#### 1.1 Normative approach for similarity-based novelty

As described in the main manuscript, we estimate the familiarity density *p*^(*t*)^ at time point *t*,

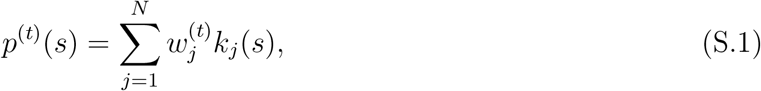

by choosing the mixture weights at time 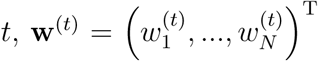 as the maximum-a-posteriori (MAP) estimate of the sequence *s*_1:*t*_ of observations up to *t*:

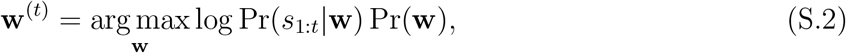

where Pr(**w**) is a Dirichlet prior over the weights, i.e.

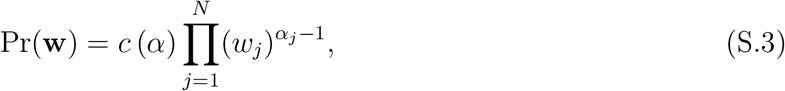

with concentration parameters *α* = (*α*_1_, …, *α*_*N*_)^*T*^ and

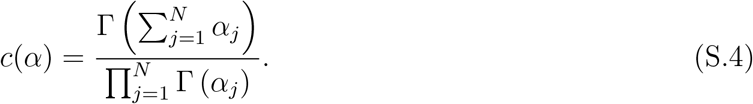

To compute the MAP estimate of the weights, we use the EM algorithm which allows us to compute the MAP estimate in an iterative fashion. The underlying idea is that, by introducing latent variables, the loglikelihood in Eq. S.2 can be decomposed into two terms that can be iteratively optimized, yielding the so-called E-step and M-step of the EM algorithm. We first derive the full EM updates for our model setup. We then show how these updates can be simplified in the incremental EM setup, a variant of EM that is more appropriate to our setting (see below for a discussion of our choice of EM algorithm).

#### 1.2 Classical EM for similarity-based novelty

##### Latent variables

To apply the EM algorithm to our mixture model, we define vectors of binary latent random variables 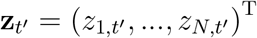 for each time step *t*′ = 1, …, *t*, such that the following three conditions hold:

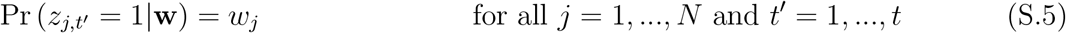

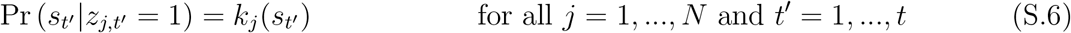

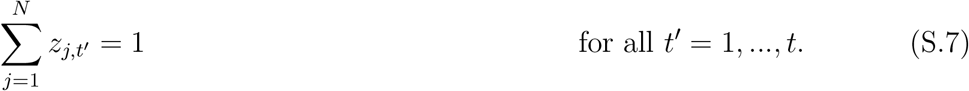

Note that the last condition, Eq. S.7, implies that each vector **z**_*t*′_ is a one-hot-coding vector of the *N* component indices. The probability of a given **z**_*t*′_ to be a one-hot-coding vector of index *j* is given by the component weight *w*_*j*_ (Eq. S.5), while the probability of observing stimulus *s*_*t*′_ when **z**_*t*′_ is encoding index *j* is the value of component *j* at stimulus *s*_*t*′_ (Eq. S.6). Effectively, the latent variables thus link the component activations for a given stimulus and the component weights.

Eq. S.5 and Eq. S.6 allow us to write conditional distributions for the stimulus and latent variable at time *t*′, *s*_*t*′_ and **z**_*t*′_:

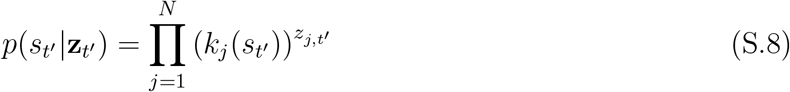

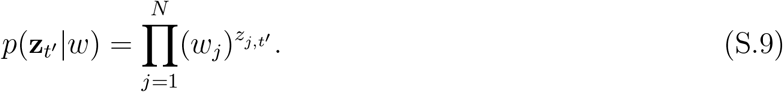

We consider the observations *s*_*t*′_ to be independent (see discussion below). That allows us to further write the joint distributions

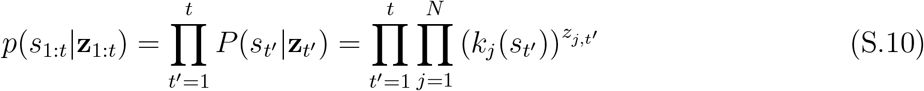

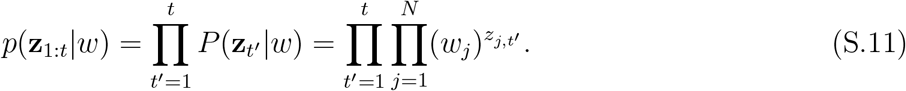

These distributions will give us the explicit expressions for the E-step and M-step that we will now derive via the loglikelihood decomposition.

##### Loglikelihood decomposition

The latent variables **z**_*t*_ allow for a decomposition of the loglikelihood in Eq. 7. Let *q*(**z**_1:*t*_) be a arbitrary distribution over the latent variables **z**_1:*t*_, then we can write the loglikelihood as

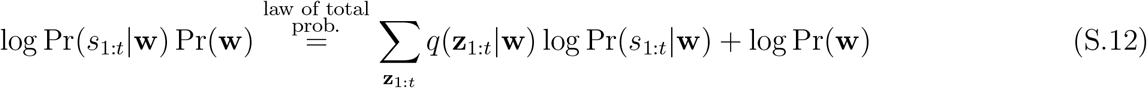

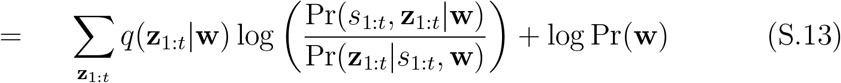

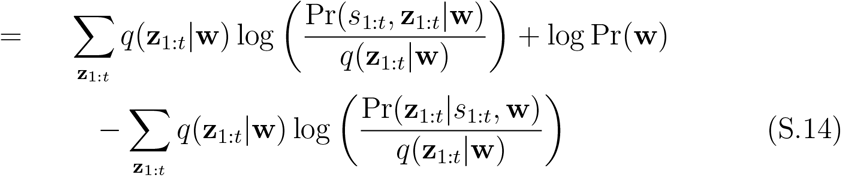

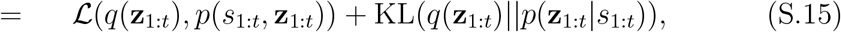

where *p*(**z**_1:*t*_ | *s*_1:*t*_) = Pr(**z**_1:*t*_| *s*_1:*t*_, **w**). Based on this decomposition, the loglikelihood can be maximized by iteratively (i) minimizing the KL divergence (E-step) while keeping the weights **w** fixed at their prior values, and (ii) maximizing the Q term with respect to **w** while keeping the latent distribution *q* fixed (M-step). For the proof that this iterative procedure indeed maximizes the loglikelihood, see e.g. Ref. [46, 71].

##### E-step

The KL-divergence of *q* and *p* becomes minimal when the two distributions are equal, so in the E-step, we set

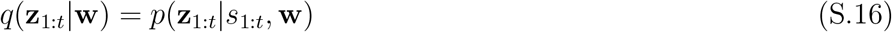

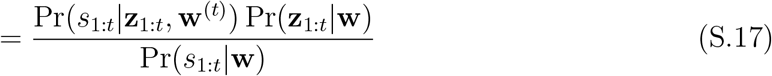

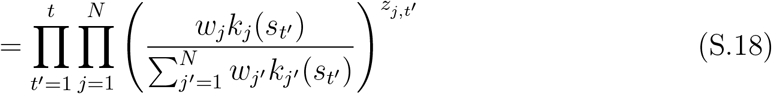

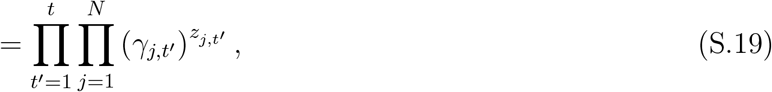

where we have used the mutual independence of any two observations *s*_*t*′_, *s*_*t*′′_ and any two latent variables **z**_*t*′_ and **z**_*t*′′_, respectively to obtain Eq. S.17, and substituted the distributions from Eq. 3, Eq. S.8 and Eq. S.9 to obtain Eq. S.18. To compute the distribution *q* in the E-step, it is thus sufficient to compute the terms

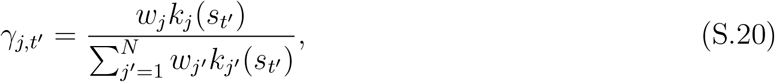

also called ‘responsibilities’, for every *j* = 1, …, *N* and every observed stimulus *s*_*t*′_, *t*′ = 1, …, *t*.

##### M-step

To maximize the term ℒ (*q*(**z**_1:*t*_), *p*(*s*_1:*t*_, **z**_1:*t*_)) with respect to **w**, we keep the latent distribution *q* fixed (denoted by fixed weights **w**_E-step_ that parametrize *q*). Note that

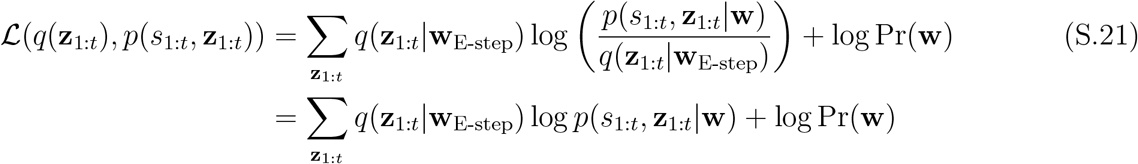

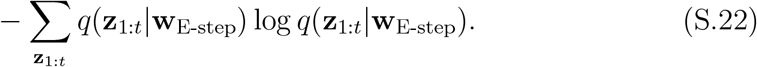

Since we keep the latent distribution *q* fixed throughout the maximization, the last term in Eq. S.22 is constant in **w**. It is thus sufficient to maximize

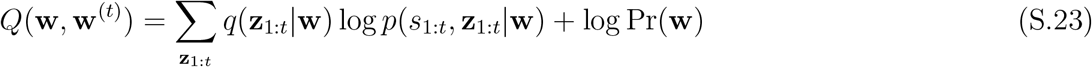

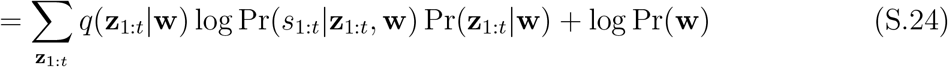

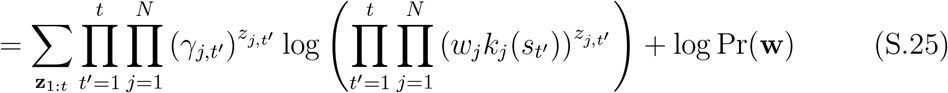

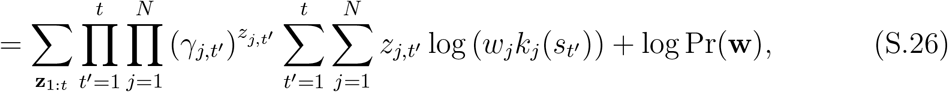

where we have substituted the distributions in Eq. S.8, Eq. S.9 and the expression for *q* from the E-step. Since all **z**_*t*′_ are one-hot coding vectors, exactly one 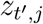 is non-zero for each *t*′; and since we are summing over all possible one-hot coding sequences of vectors **z**_1:*t*_, each *z*_*j*,*t*′_ will be non-zero exactly once. Eq. S.26 therefore reduces to

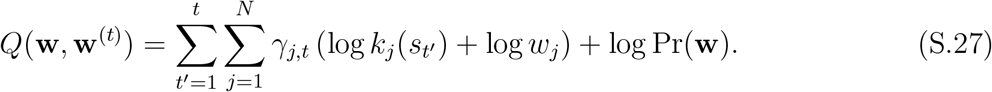

We maximize *Q* as in Eq. S.27 under the constraint that all weights sum to one using Lagrange multipliers, by solving

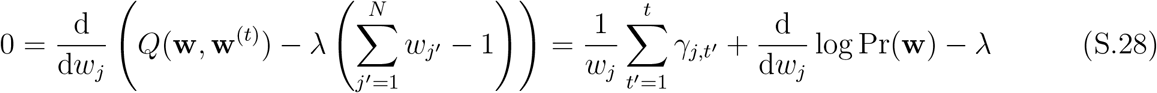

for all *j* = 1, …, *N*. Since the Lagrange multiplier has to satisfy Eq. S.28 for all *j*, it has to satisfy the following equation (after multiplying Eq. S.28 with *w*_*j*_ and summing over *j*):

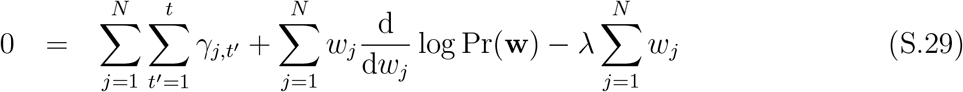

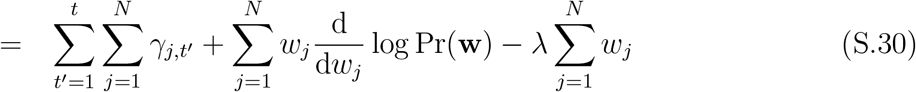

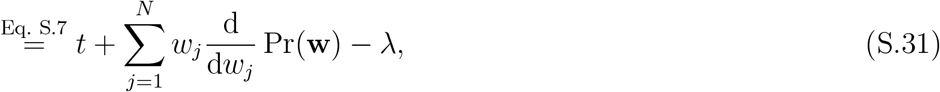

yielding

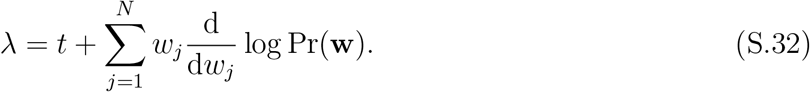

Substitution in Eq. S.28 gives the solution for the new mixture weights:

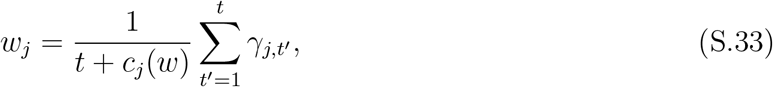

where

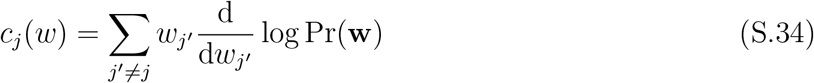

##### Special Dirichlet priors

For our Dirichlet prior (Eq. S.3), *c*_*j*_ (*w*) reduces to

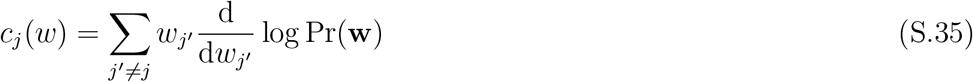

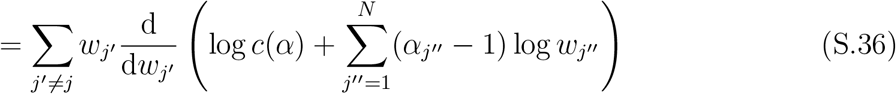

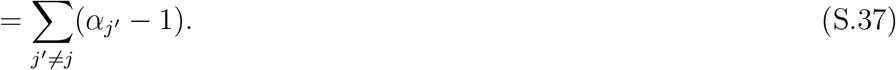

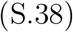

For Dirichlet prior with shared concentration parameter *α*_*j*_ = *ϵ* + 1 for all *j* = 1, …, *N*, *c*_*j*_ (*w*) reduces to (*N* − 1)*ϵ* for all *j*. In this case, the M-step update (Eq. S.33) reduces to

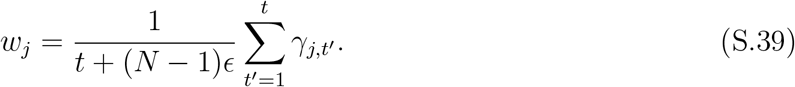

A Dirichlet prior with shared concentration parameter encodes the assumption that all components have been observed equally prior to observing the stimulus sequence *s*_1:*t*_ – a reasonable assumption that we will make for the rest of the derivation.

#### 1.3 Incremental EM for similarity-based novelty

##### Full EM

In summary, the full EM algorithm estimates the mixture weights that maximize the likelihood of our observations *s*_1:*t*_ in two alternating steps: first, the ‘responsibilities’ *γ*_*j*,*t*′_ are updated for all *j* = 1, …, *N* and *t*′ = 1, …, *t* as in Eq. S.20 (E-step); then the weights 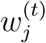 after observations *s*_1:*t*_ are computed as in Eq. S.39 (M-step). In the full EM setup, these two update steps are repeated until convergence of the algorithm. Since the full EM relies on knowledge about all observations *s*_1:*t*_ to make a single update, the mixture weights can only be estimated after observing the full sequence *s*_1:*t*_ that is then processed together in every iteration of the EM algorithm (‘batch mode’).

##### Incremental EM (I-EM)

For our goal of modeling human and animals, however, this ‘batch’ update of the familiarity density (and hence, of stimulus novelty) is undesired since we assume that the brain updates its estimate of the novelty of a stimulus immediately after its observation – even if that leads to a less precise estimate of the overall statistics of stimuli in the environment. Instead of the full EM algorithm, we therefore estimate our component weights using the incremental EM algorithm. This variant of EM that uses only a single observation in each update iteration, iterating through the sequence of observations *s*_1:*t*_ one-by-one, and then starting again at the beginning of the sequence until the algorithm has converged.

##### Continual incremental EM (CI-EM)

For our scenario of humans and animals experiencing an unfamiliar environment, we assume that the incremental EM algorithms uses a continuing sequence of observations for its estimates, such that each observation is used only once for the update of the algorithm. The resulting weight estimates are in that sense ‘approximate’ estimates of the component weights but also the best estimates that are available for a potentially infinite sequence of observations. Moreover, in environments with finitely many distinguishable stimuli, all stimuli will eventually be revisited and thus contribute again to the weight estimate (like in the incremental EM for finite observation sequences). An important difference is, thought, that the proportion of how much each stimulus is revisited and thus contributes to the estimate of the familiarity distribution is not determined by a fixed circular schedule but by the experimenter (passive viewing task) or by the human or animal through its action policy (active exploration). To distinguish our specific use of the incremental EM algorithm from its classical ‘finite stimulus sequence’ usage, we also refer to our algorithm as ‘continual incremental EM algorithm’.

##### Continual incremental EM (CI-EM) updates

The main difference between full EM and incremental EM updates is that the latter only uses a single observation *s*_*t*′_ in its E-step update: instead of recomputing the responsibilities for all latent variables *z*_*j*,*t*′′_, for all *j* = 1, …, *N* and all *t*^*′′*^ = 1, …, *t*, the incremental E-step only recomputes the responsibilities for the observation *s*_*t*′_; all responsibilities from previous iterations of the algorithm are kept the same:

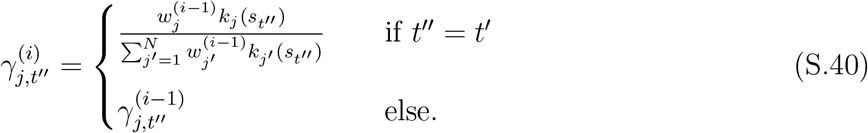

where the additional index *i* denotes the *ith* iteration of the algorithm, in which observation *s*_*t*′_ is considered. The M-step of the algorithm is the same as in the full EM algorithm (Eq. S.39) – just the underlying responsibilities are different due to the incremental E-step.

In our continual I-EM algorithm, each observation *s*_*t*_ is followed by one iteration of the I-EM algorithm, i.e. the *t*-th step of algorithm considers the *t*-th observation:

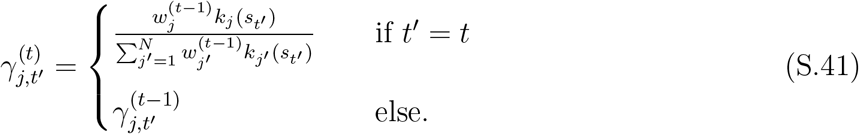

In this scenario, the *t*-th M-step update (Eq. S.39) can be written as

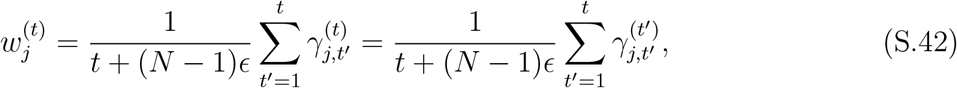

where 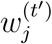 for *j* = 1, …, *N* are the component weights at time *t*′ in the experiment. Eq. S.42 gives rise to an iterative update rule (also see Methods, where we dropped the iteration index *t*′ for better readability):

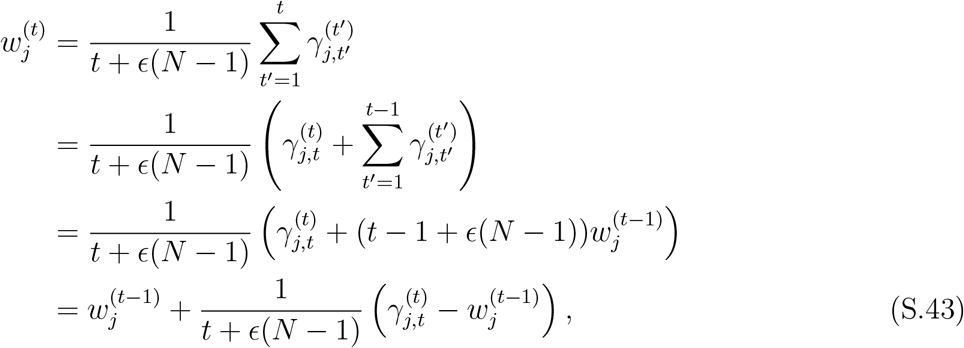

where 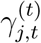 is given by Eq. S.41.

#### 1.4 Non-leaky count-based novelty as a special case of similarity-based novelty

In a scenario where all observations are (perfectly) distinct, count-based novelty and similarity-based novelty are equivalent. To formalize this statement, consider a sequence of observations *s*_1:*t*_, where each observation is drawn out of a set of *N* perfectly distinct stimuli, *S* = {*s*_1_, …, *S*_*N*_}. We say that count-based novelty and similarity-based novelty are equivalent with respect to this sequence if, at all time steps *t*′ = 0, …, *t* and for all stimuli *s*_*j*_ ∈*S*, their familiarity distributions 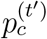 (count-based) and 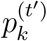 (similarity-based) take identical values, i.e.

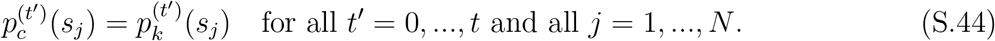

We now show that Eq. S.44 holds for count-based and similarity-based novelty. To this end, first note that the count-based familiarity at each time point *t* is given as

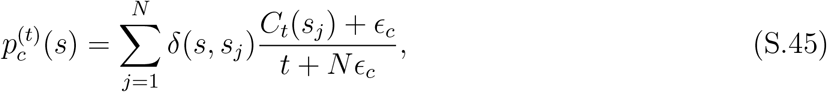

where *ϵ*_*c*_ is the count-based prior and *δ* is the Kronecker delta function. We can rewrite the count-based familiarity distribution (Eq. S.45) in the style of a similarity-based familiarity density (i.e. as a mixture model) as follows:

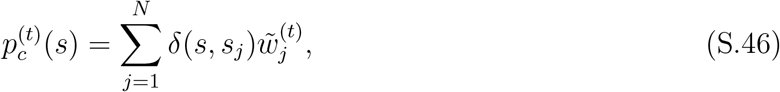

with

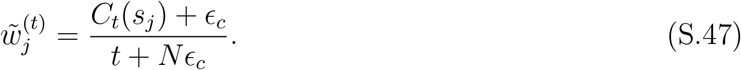

Since the counts are initialized as *C*_0_(*s*_*j*_) = 0 for all *j* = 1, …, *N*, the initial ‘count-based’ mixture weights at time *t* = 0 are given as

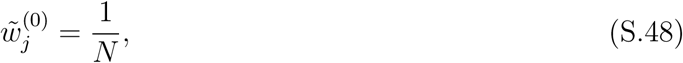

for all stimuli *j* = 1, …, *N*. Since counts are updated at subsequent time steps *t* as

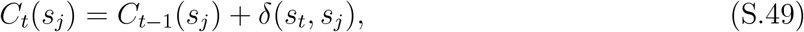

the update of the count-based mixture weights is given as:

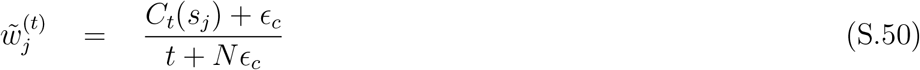

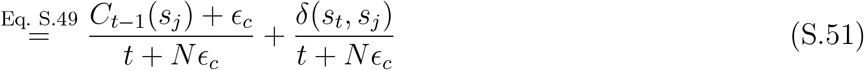

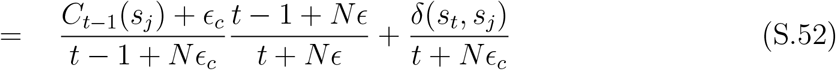

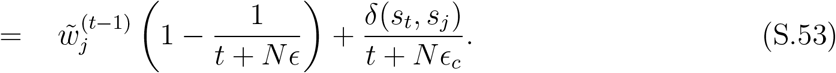

We now construct a similarity-based novelty model that is equivalent to the previous count-based novelty model. Since all observations *s*_*j*_ are perfectly distinct, we can construct a continuous stimulus space *S*′ ⊃ *S* and non-overlapping, box-shaped components *k*_*j*_, *j* = 1, …, *N*, such that

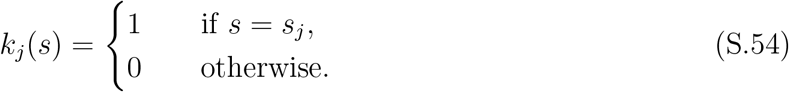

and 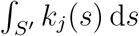 for all *j* = 1, …, *N* (e.g., we can place the *s*_*j*_ on a real line with equal distance 1 between them, and define box components of height and width 1, centered at the different *s*_*j*_). Then the familiarity distribution of the similarity-based novelty model at stimulus *s*_*i*_ ∈*S* is given as

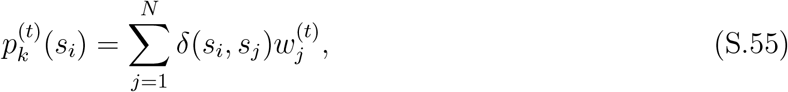

By comparing with Eq. S.46, we can see that, for the equality in Eq S.44 to hold, it is sufficient to show that 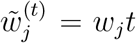 for all *j* and all *t*. The component weights are uniformly initialized to 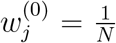, and thus equal to the count-based initial weights (Eq. S.48). The component weights are updated with the incremental update rule derived above (Eq. S.43), i.e.

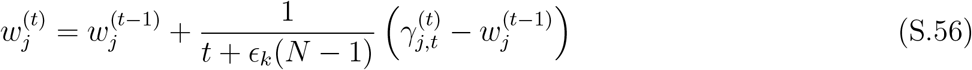

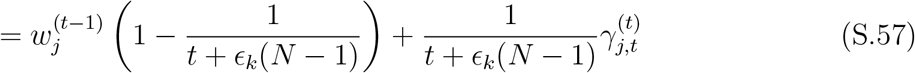

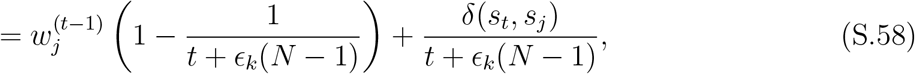

where we have used that, given our choice of box components, we have

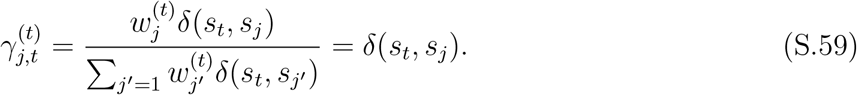

By comparing with the update rule of the count-based mixture weights (Eq. S.53), we can see that by setting the similarity-based novelty prior 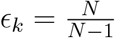, the two weight updates become equivalent, and, hence, the two novelty definitions are equal in the sense of Eq. S.44.

#### 1.5 Network implementation of similarity-based novelty

**Figure 5:**
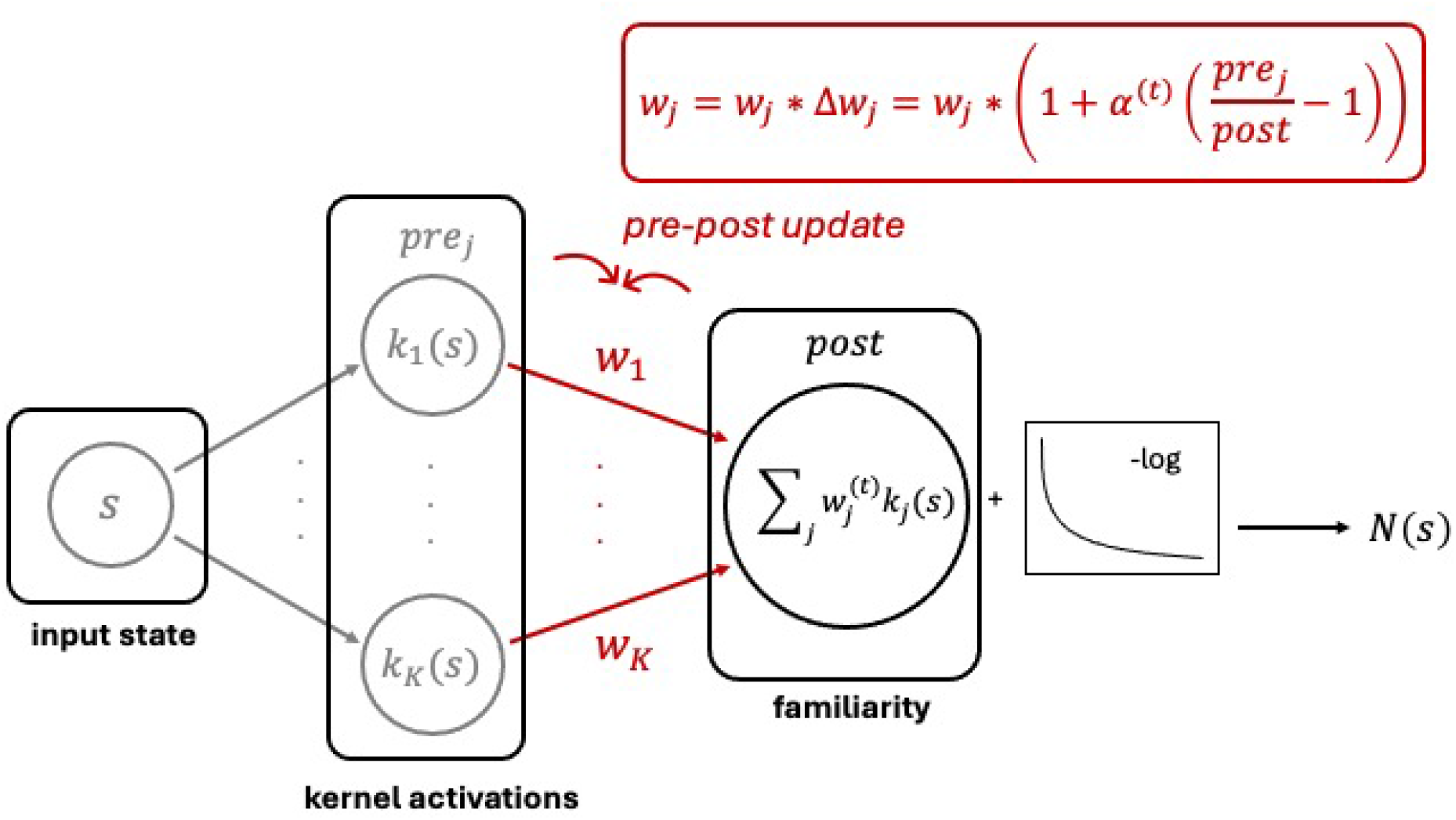
Network implementation of similarity-based novelty. The weight update rule of similarity-based novelty (Eqs. 4,5 in main text) can be written as a multiplicative weight update rule (red box) in a biologically realistic circuit. The amplitude of weight updates depends on the ratio of presynaptic activity (component activation by current input stimulus *s*) and postsynaptic activity (familiarity of current input stimulus *s*, given the previous weights).

## 2 Passive viewing experiment: data set and modeling

### 2.1 Extraction of neural responses in Homann et al

Population responses are extracted from the raw fluorescence traces in six steps: (i) fluorescence traces *F* (*t*) for each ROI (manually identified) are obtained by averaging over the ROI’s pixels on a frame-by-frame basis; (ii) relative fluorescence traces for each ROI are computed as 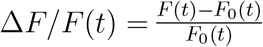, where a smooth estimate of the baseline fluorescence *F*_0_(*t*) is computed using spline interpolation; (iii) event-triggered activity (ETA) traces for each event type (novel image, familiar image, recovery probe) is computed by trial-averaging relative fluorescence traces Δ*F/F* (*t*) in the window 1000 ms before until 300-500 ms after the event, separately for each ROI; (iv) excess activity traces in each ROI for the novel image and the recovery probe events are obtained by subtracting the ETA for familiar image events from the respective ETAs of novel image and recovery probe events; (v) population excess activity traces (and population steady-state activity traces) are computed as the population average over the individual ROI’s excess (and steady-state) activity traces; (vi) the population’s novelty and recovery probe responses Δ*N* are given by the maximum amplitude of the population excess activity to the respective event (novel image or recovery probe), while the population’s steady-state response *N*_∞_ to familiar images is calculated as the average of the population steady-state activity trace.

### 2.2 Grid search: parameter stability

**Figure 6:**
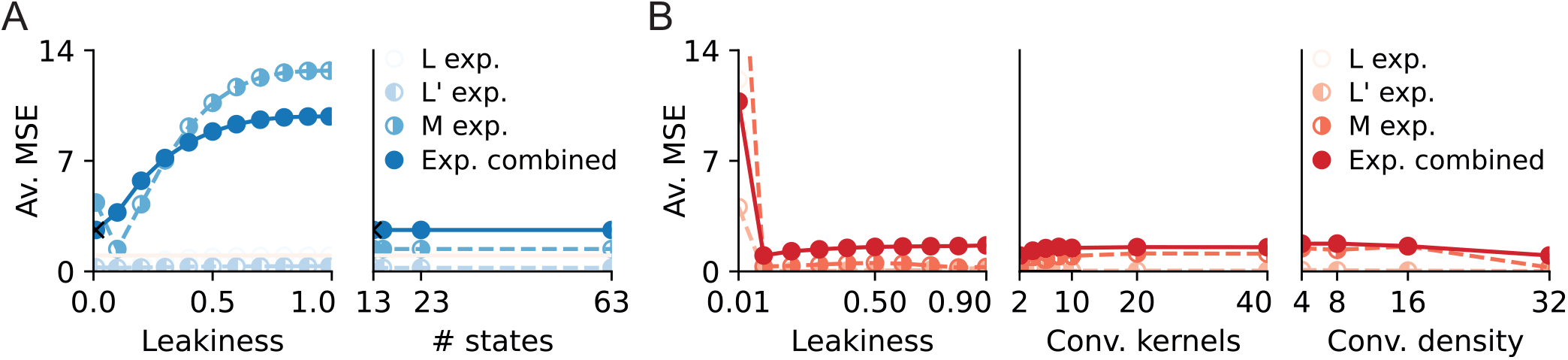
Robustness of fit to neural data. Individual MSE for each experiment by Homann et al. [38] and average MSE across experiments for different values of (A) the count-based novelty parameters *α* (leakiness) and |*S*| (number of states; and (B) the similarity-based novelty parameters *α* (leakiness), *c* (number of different features encoded by components) and *d* (number of convolutional components for each feature).

### 2.3 Linear regression fit to neural data

For each parameter set *θ* of a given model ℳ, we use least-squares estimation (LSE) to find the regression coefficients *β* that minimize the regression error *ϵ* in the regression equation

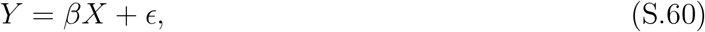

where *Y* is the neural data vector, and *X* contains the model predictions for *Y*. The data vector *Y*∈ ℝ^20^ contains the average neural responses to novelty (and familiarity) measured in the three experiment variations:

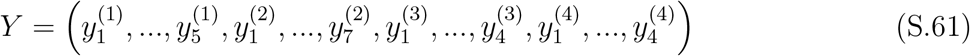

where 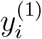 are the average neural novelty responses Δ*N* during the ‘variable repetition experiment’ for different number of repetitions *L* of the familiar sequence 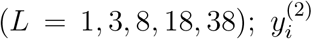 are the average neural novelty responses Δ*N* to the formerly familiar sequence in the ‘repeated image set experiment’, for different lengths Δ*T* of the replacement interval (Δ*T* = 0, 21, 42, 63, 84, 108, 144 seconds); and 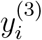 and 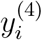 are the average neural responses to novelty, Δ*N*, and to familiarity, *N*_∞_, respectively, during the ‘variable image number experiment’ for different number *M* of familiar stimuli (*M* = 3, 6, 9, 12). The model predictions *X* ∈ ℝ^20×2^ are given as

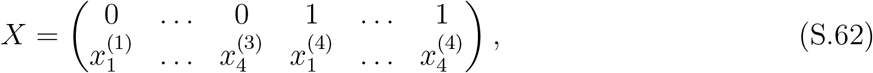

where the row column, containing the model’s respective predictions for the novelty and familiarity signals in each of the experimental conditions underlying the data vector *Y*, is used to fit the scaling factor *β*_2_ (second entry of *β* ∈ ℝ^2^) of the linear regression, and the first row is used to fit the shift *β*_1_ (first entry of *β* ∈ ℝ^2^) of the predicted steady-state responses *N*_∞_ relative to the actual neural signals to familiar stimuli. Note that since the novelty responses Δ*N* are normalized with the steady-state responses, the shift does not apply to them, such that they have zero-entries in the first row of *X*.

For each model ℳ, we choose the parameter set *θ* that minimizes the mean squared error (MSE) between neural data *Y* and the fitted model predictions *βX*^*^, normalized with the variance of the neural data *σ*^2^(*Y*):

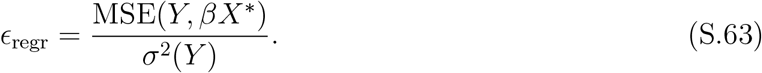

## 3 Active exploration experiment: data set and modeling

### 3.1 Behavioral data by Rosenberg et al.: Outliers

For mouse ‘B2’, which fails to receive a reward during its first 6 visits to the goal state, we only consider behavior up to its first goal encounter. Like Rosenberg et al., we exclude mouse ‘D6’ since it fails to enter the maze or more than one short (1 time step) bout. In contrast to Rosenberg et al., we include mouse ‘C6’ in our analysis since its behavior with respect to our behavioral statistic does not show outlier behavior. While Rosenberg et al. analyze exploration behavior based on ‘first water runs’, which includes the direct paths between maze entry and the goal state in the maze, we analyze the entire state trajectory of mice during the exploration between first entry and encounter of the goal state.

### 3.2 Novelty-seeking RL (N-RL) algorithms

#### 3.2.1 Model-based N-RL algorithm

The model-based (MB) N-RL algorithm (Alg. 1) uses prioritized sweeping (e.g., see [74]) for the model-based update of the novelty-based Q-values (‘N-Q-values’), and applies a softmax policy on the N-Q-values to determine its actions.

The MB N-RL algorithm is characterized by the parameters of the novelty model (*ϵ*_*c*_ for count-based novelty; *ϵ*_*k*_ and **k** for similarity-based novelty, see Normative update rule for similarity-based novelty), as well as five additional parameters (see line 2 in Alg. 1). The inverse temperature *β* determines the noise level of the agent’s softmax policy over the Q-values (low *β* leads to more noisy actions). The leak factor *k*_leak_ and the prior *ϵ*_env_ of the belief counts *α* determine how strongly the *α* values for a given transition decay between two observations of that transition – the higher *k*_leak_, the more influenced are the *α* values by recently observed transitions, and the lower the influence of the prior *ϵ*_env_. A lower *ϵ*_env_, in turn, causes the *α* values to adjust more quickly to the observed transition counts. The more deterministic our environment is, the lower our *ϵ*_env_ should thus be, since every observation contains reliable information about the underlying state transition. The parameter *λ* and *T*_*PS*_ characterize the Q-value updates: *T*_*PS*_ determines how many sweeps, i.e. Q-value updates are performed in each step (the higher *T*_*PS*_, the more precise the model-based Q-values update), while *λ* has a similar role to the discount factor *γ* that discounts the influence of future expected novelty on the state value of a given state *s* (the higher *λ*, the more important the value of future novelty).

#### 3.2.2 Model-free N-RL algorithm

The model-free (MF) N-RL algorithm (Alg. 3) implements an actor-critic algorithm [74] that updates novelty-based V-values using TD-learning and novelty-based action preferences using policy gradient learning. Actions are chosen using a softmax policy over the action preferences. In addition to the parameters of the novelty model (same as above for the MB N-RL algorithm), the MF N-RL algorithm is characterized by 8 parameters: the inverse temperature *β* of the softmax action policy, the initial V-values *V*_0_ and action preferences *h*_0_, the learning rates for critic (*α*_*c*_) and actor (*α*_*a*_), the discount factor for future novelty *γ*, and the two decay factors of the critic and actor eligibility traces *λ* and 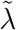.

##### Algorithm 1 Model-based N-RL algorithm

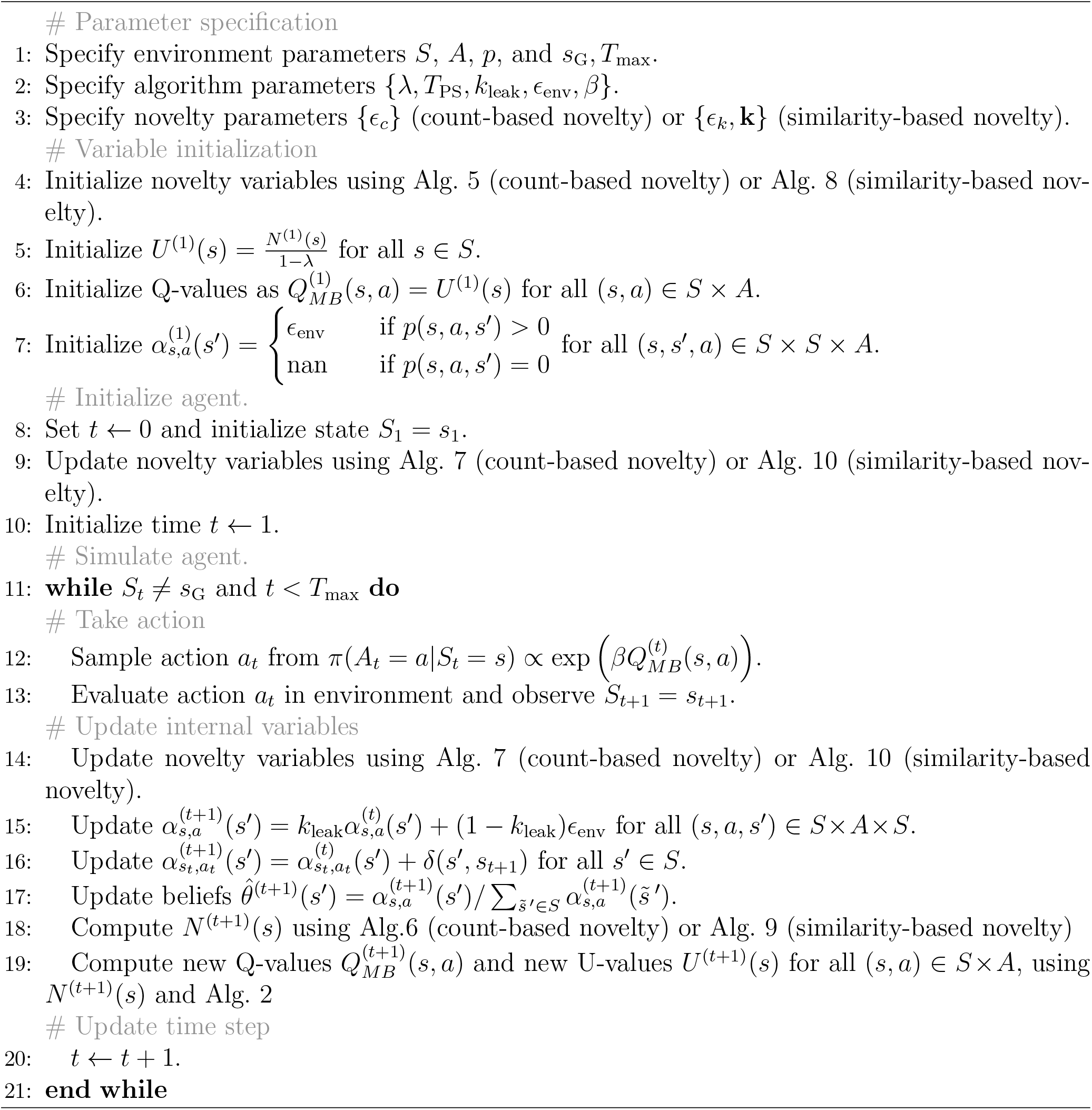

#### 3.2.3 Hybrid N-RL algorithm

The hybrid N-RL algorithm runs the model-based and the model-free N-RL updates in parallel and uses a hybrid policy that is based on a linear combination of the model-based novelty Q-values and the model-free novelty action preferences.

The hybrid N-RL model has one set of novelty parameters, analogously to the MF and MB N-RL algorithms above, as well as the respective MF and MB algorithm parameters from Algs. 1,3. The only additional parameter is the weight *w*_*MF*_ that determines the balance of the MF and MB softmax distribution in the hybrid policy.

##### Algorithm 2 Prioritized Sweeping with target signal *N* ^(*t*+1)^

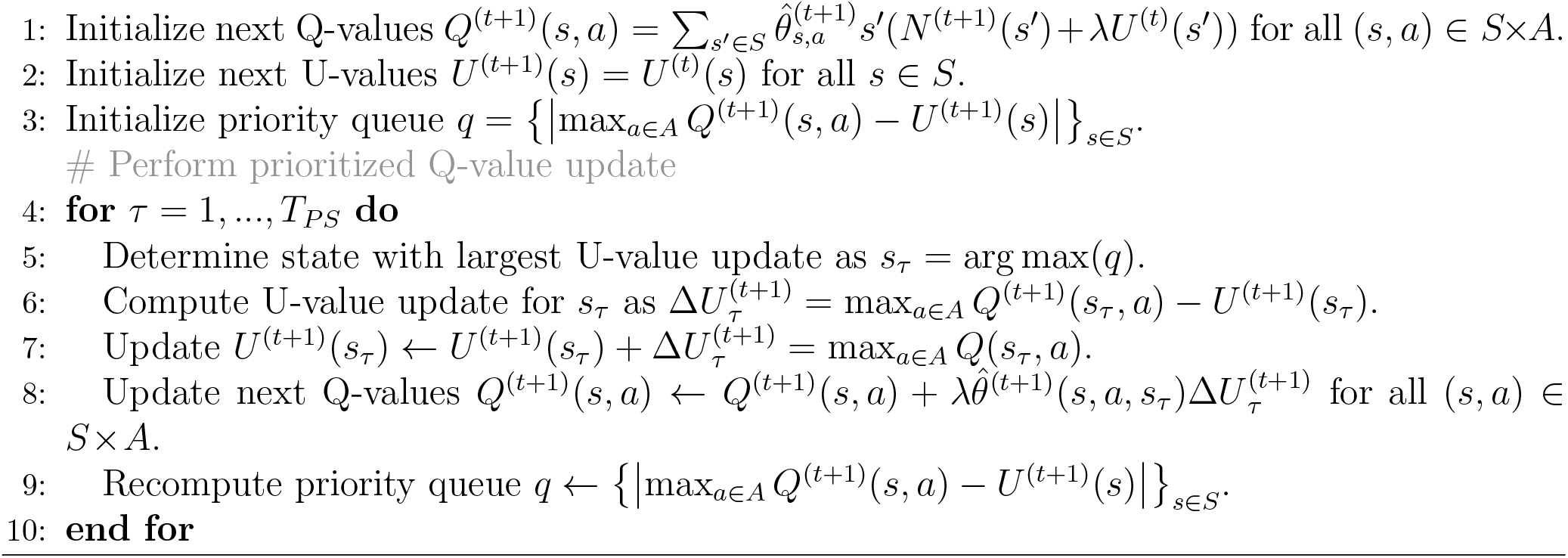

##### Algorithm 3 Model-free N-RL algorithm

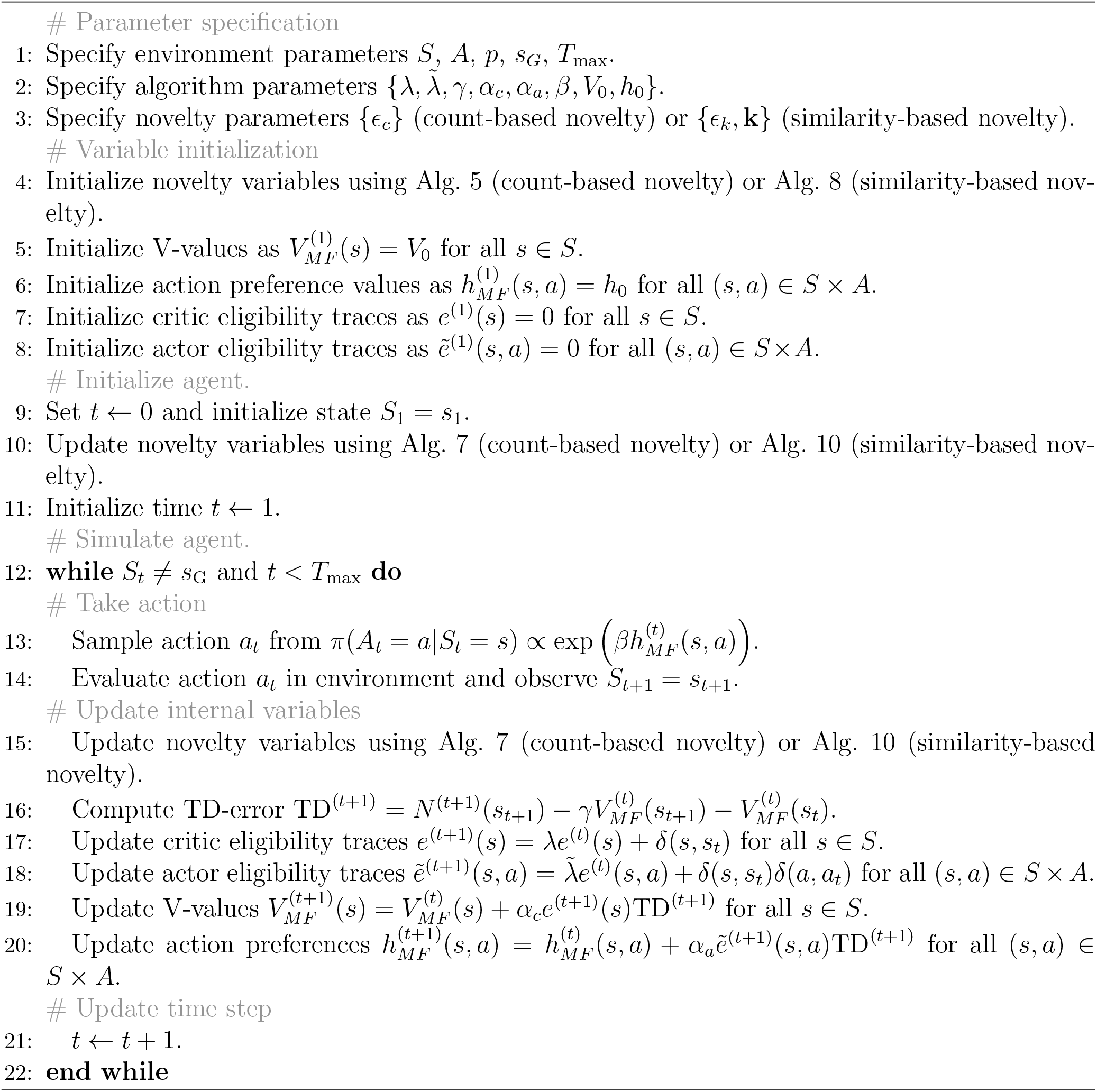

##### Algorithm 4 Hybrid N-RL algorithm

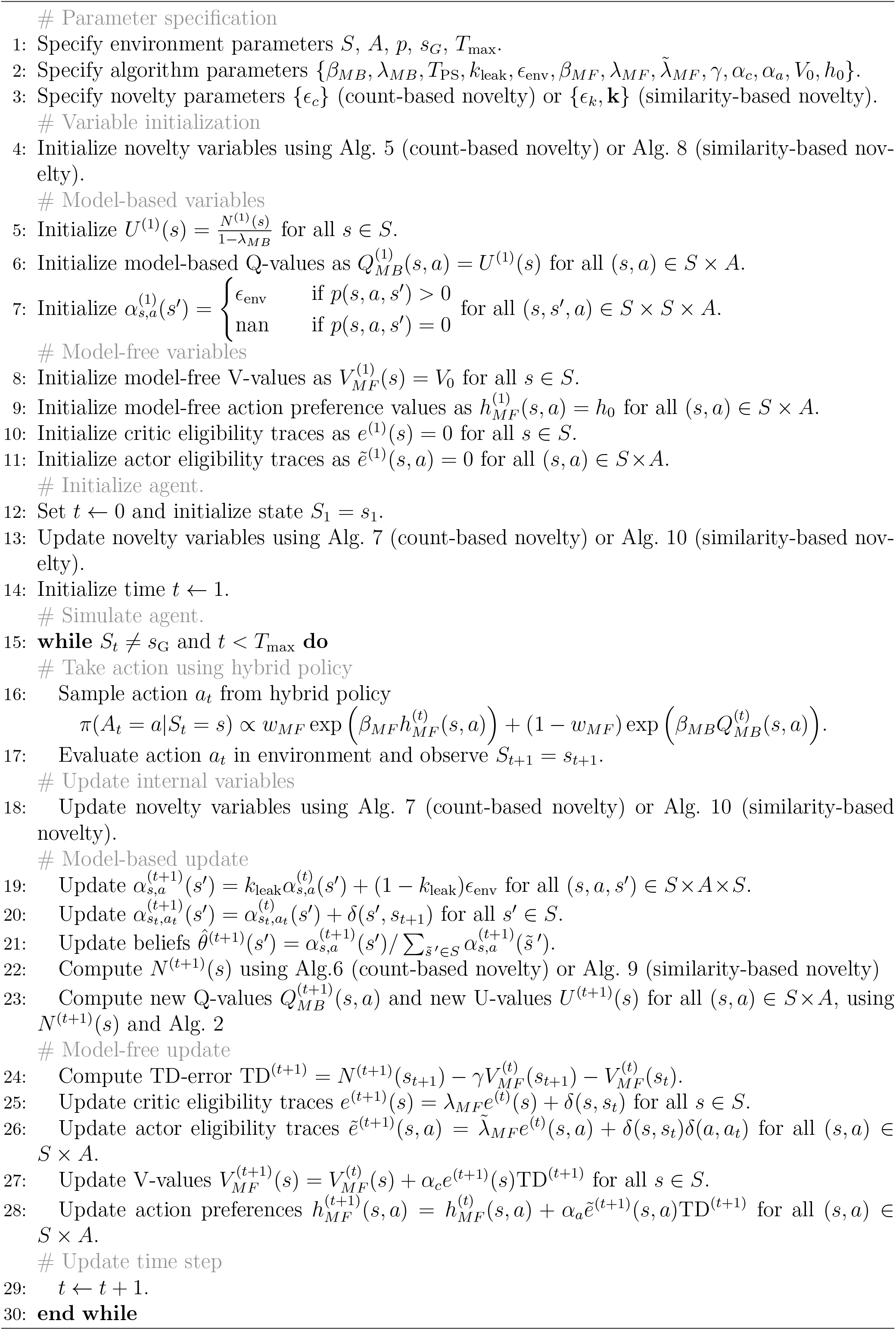

### 3.3 Novelty models for N-RL algorithm

#### Algorithm 5 Novelty initialization (count-based novelty)

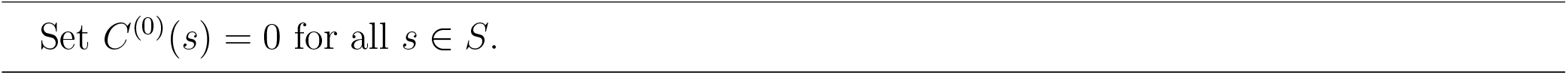

#### Algorithm 6 Novelty evaluation (count-based novelty)

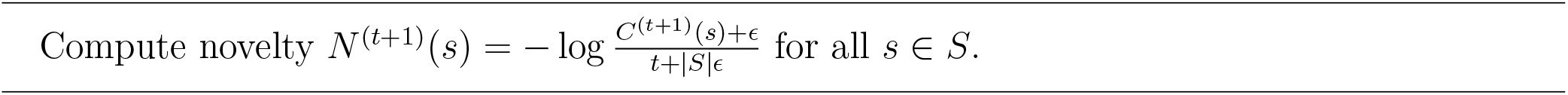

#### Algorithm 7 Novelty update (count-based novelty)

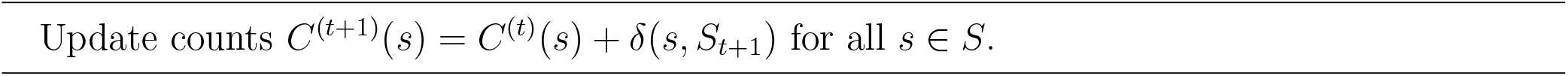

#### Algorithm 8 Novelty initialization (similarity-based novelty)

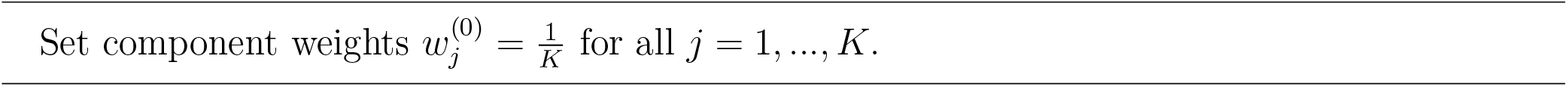

#### Algorithm 9 Novelty evaluation (similarity-based novelty)

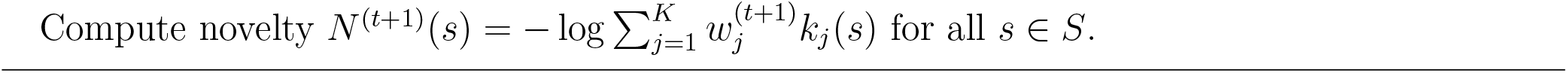

#### Algorithm 10 Novelty update (similarity-based novelty)

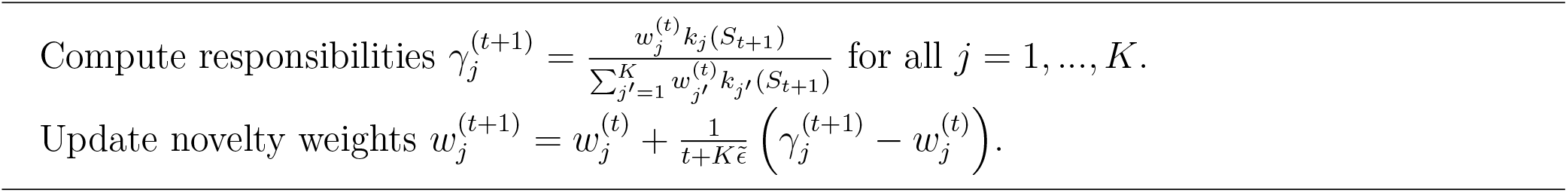

### 3.4 Behavioral fit: MLE derivation for N-RL models

To compute the log-likelihood in Eq. 24 for a given N-RL model, we rewrite it in terms of the model’s softmax policy *π*(·|ℳ, *θ*) (see SI) as

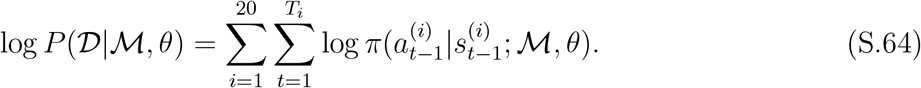

To compute the right-hand side of Eq. S.64, we simulate the model ℳ with parameters *θ* for each mouse trajectory; but, instead of letting the model choose the action in each step *t* according to its policy, we choose the corresponding action 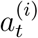 from the mouse data and compute its log-likelihood under the model’s policy. These individual log-likelihoods are then added to obtain the total likelihood of the data under the given model ℳ with parameters *θ* (Eq. S.64).

The expression for the log-likelihood in Eq. S.64 can be derived as follows:

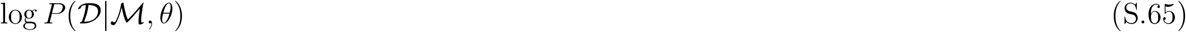

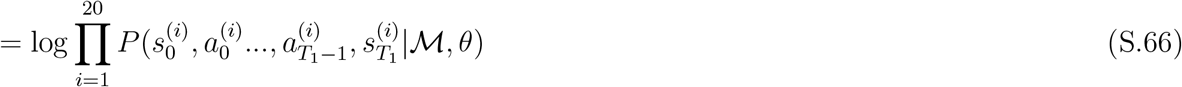

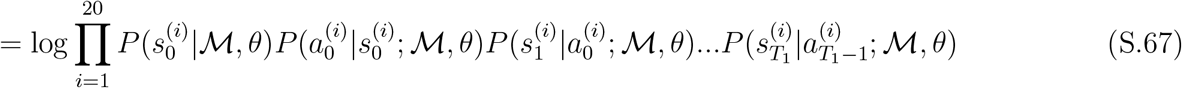

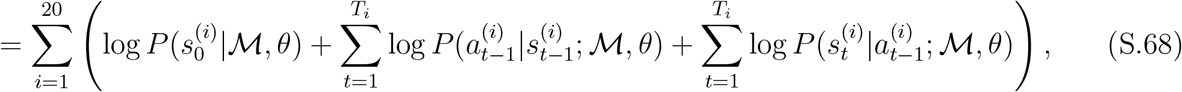

where we have use the Markov property of the N-RL models (see above). Note that 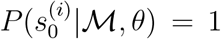 since all mice and agents start in the home cage by experiment design; and that 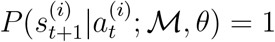 since the environment is deterministic, i.e. given action *a* always leads to the same state *s*. Using this, we can simplify Eq. S.68 to

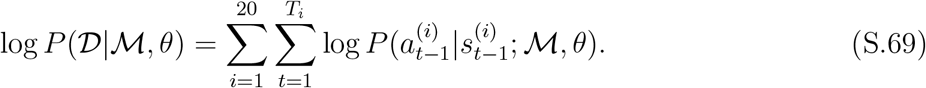

We can further write the individual log-likelihoods in Eq. S.69 in terms of the model’s action policy *π*(·|ℳ, *θ*) (in the case of our N-RL models, a softmax policy over the model’s Q-values):

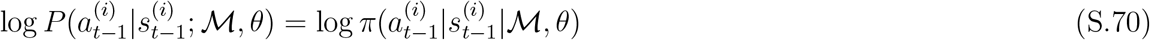

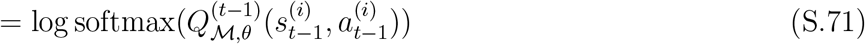

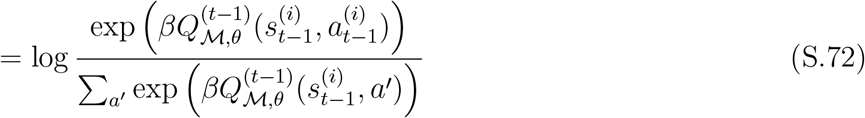

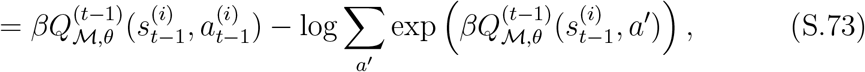

where the inverse temperature *β* of the softmax policy is part of the set of model parameters *θ*.

### 3.5 Behavioral fit: Model recovery

**Figure 7:**
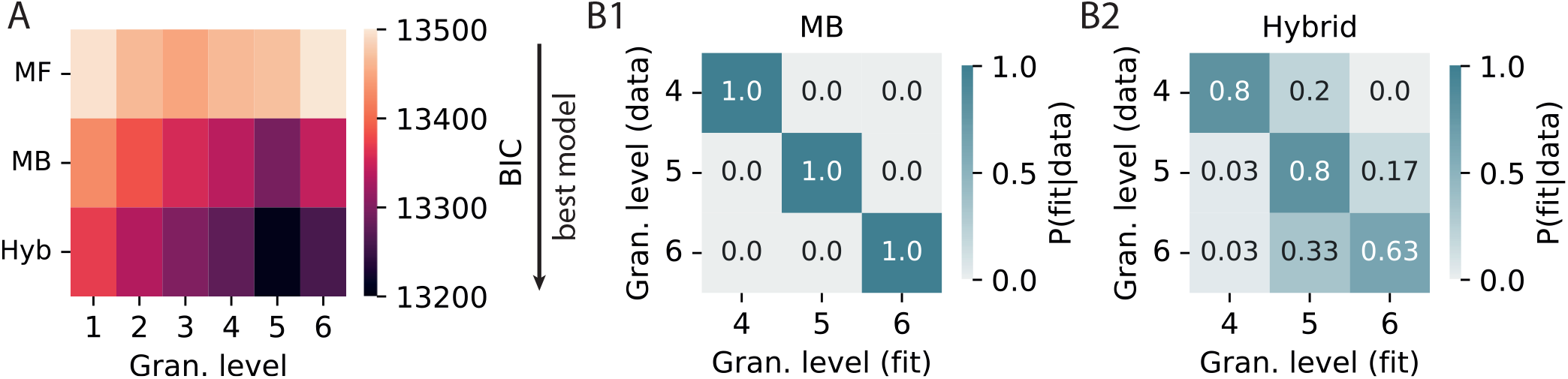
Model recovery for all similarity-based novelty models in the Rosenberg task.

## Notes

### Competing Interest Statement

The authors have declared no competing interest.

### Summary of Updates

Results section and related methods sections were updated with additional analyses.

